# Integrating Hi-C and FISH data for modeling 3D organizations of chromosomes

**DOI:** 10.1101/318493

**Authors:** Ahmed Abbas, Xuan He, Bin Zhou, Guangxiang Zhu, Zishan Ma, Jun-Tao Gao, Michael Q Zhang, Jianyang Zeng

**Affiliations:** Institute for Interdisciplinary Information Sciences, Tsinghua University, Beijing 100084, China.; School of Life Science, Tsinghua University, Beijing 100084, China.; MOE Key Laboratory of Bioinformatics; Bioinformatics Division, Center for Synthetic and Systems Biology, BNRist; Department of Automation, Tsinghua University; Center for Synthetic and Systems Biology, Tsinghua University, Beijing 100084, China.; Department of Biological Sciences, Center for Systems Biology, the University of Texas at Dallas, Richardson, TX 75080-3021, USA.; Department of Basic Medical Sciences, School of Medicine, Tsinghua University, Beijing 100084, China.

## Abstract

The new advances in various experimental techniques that provide complementary information about the spatial conformations of chromosomes have inspired researchers to develop computational methods to fully exploit the merits of individual data sources and combine them to improve the modeling of chromosome structure. In this paper, we propose GEM-FISH, a first method for reconstructing the 3D models of chromosomes through systematically integrating both Hi-C and FISH data with the prior biophysical knowledge of a polymer model. Comprehensive tests on a set of chromosomes for which both Hi-C and FISH data were available have demonstrated that GEM-FISH can reconstruct the 3D models of chromosomes with more accurate spatial organizations of TADs and compartments than using only Hi-C data. In addition, GEM-FISH can accurately capture the spatial proximity of loop loci and the colocalization of loci from the same sub-compartments. Moreover, our reconstructed 3D models of chromosomes revealed novel patterns of spatial distributions of super-enhancers which can provide useful insights into understanding the functional roles of these super-enhancers in gene regulation. All these results demonstrated that, through integrating both Hi-C and FISH data into a unified framework, GEM-FISH can provide a better tool for modeling the 3D organizations of chromosomes than using the Hi-C data alone.

## 1 Introduction

Determining the three-dimensional (3D) structure of a chromosome can provide important mechanistic insights into understanding the underlying mechanisms of the 3D folding of the genome and the functional roles of high-order chromatin compaction in gene regulation. For instance, the 3D organization of a chromosome and the spatial proximity of genomic loci can reveal essential relationships between functional elements and their distal targets along the genome sequence, which can shed light on their regulatory functions in controlling gene activities. Recently, the chromosome conformation capture (3C) technique [1], which measures the interaction frequencies between pairs of genomic loci through a proximity ligation strategy, has significantly advanced the studies of higher-order chromatin structure. The extended 3C techniques, such as Hi-C [2] and ChIA-PET [3], have enabled one to study the genome-wide landscape of 3D genome structure at different resolutions (i.e., ranging from Mbps to Kbps) and in various cell types, organisms, and conditions.

Based on their proposed Hi-C technique, Liberman-Aiden et al. [2] discovered that chromosomes are generally partitioned into two compartments, i.e., A and B, which are enriched with active and inactive chromatin marks, respectively. Using Hi-C maps with a resolution in the order of tens of Kbps, Dixon et al. [4], introduced the concept of topologically associated domains (TADs), which are defined as the regions that have higher contact frequencies within a domain than across different domains in the Hi-C maps. With Hi-C maps of a relatively high resolution (in a range of 1-5 Kbps), Rao et al. [5] were able to study a finer scale of chromatin structure and investigate the formation of chromatin loops.

Despite the recent significant progress in the studies of higher-order architecture of the genome using the 3C-based techniques, our current understanding on the 3D packing of chromosomes still remains largely incomplete. For example, there still exists a gap in understanding the spatial organizations of the A/B compartments relative to each other in individual chromosomes. Also, if two genomic loci have relatively low contact frequency, it is usually difficult to infer their relative spatial positions only from the Hi-C maps. On the other hand, the fluorescent in situ hybridization (FISH) technique, which measures the spatial distances between a pair of distal genomic loci over a number of cells through a direct imaging strategy, can provide a complementary tool to investigate the 3D organizations of chromosomes.

Wang et al. [6] applied a multiplexed FISH method to study the spatial organizations of TADs and compartments in Chromosomes 20, 21, 22, and X of human diploid (XX) IMR90 cells. They observed that the relation between spatial and genomic distances might deviate from the 1/3 power law expected from the ideal fractal globule model [7], especially when the genomic distance exceeds 7 Mbps. In addition, they found that the A/B compartments are usually arranged in a spatially polarized manner relative to each other with different compartmentalization schemes for the active (ChrXa) and inactive (ChrXi) states of X-Chromosomes. In particular, for ChrXi, the A/B compartments are separated by the DXZ4 macrosatellite, while for ChrXa, these two compartments correspond to the p and q arms.

A large number of computational methods have been developed in the past few years to determine the 3D structures of chromosomes from Hi-C maps [1,8–31]. Many of these methods estimate the pairwise spatial distances between genomic loci using the formula *f* ∝ 1/*d^α^*, where *f* and *d* stand for the contact frequency and the estimated spatial distance between a pair of loci, respectively, and *α* is a constant. Recently, our group has developed a new manifold learning based approach, called GEM [32], which combines both Hi-C data and conformational energy derived from our current available biophysical knowledge about a 3D polymer model to calculate the 3D structure of a chromosome. GEM does not depend on any specific assumption about the relation between the Hi-C contact frequencies and the corresponding spatial distances, and directly embeds the neighboring proximity from Hi-C space to 3D Euclidean space. Comprehensive comparison tests have demonstrated that GEM can achieve better performance in modeling the 3D structures of chromosomes than other state-of-the-art methods [32].

Despite the recent new advances in FISH techniques [33–36], obtaining a high-resolution pairwise distance map similar to a Hi-C contact map in the same high-throughput manner is still out of reach [37]. On the other hand, the large amount of available FISH data provide an important source of complementary constraints to Hi-C maps for modeling the 3D architectures of chromosomes. However, integrating both Hi-C and FISH data into a unified framework for modeling 3D chromosome structures is not a trivial task, and requires the development of a systematic data integration approach to fully exploit the strengths of individual data types to improve the modeling accuracy. To our best knowledge, no computational approach has been proposed previously to integrate both Hi-C and FISH data for reconstructing the 3D models of chromosomes.

In this paper, we propose a novel divide-and-conquer based method, called GEM-FISH, which is an extended version of GEM [32] and the first attempt to systematically integrate FISH data with both Hi-C data and the prior biophysical knowledge of a polymer model to reconstruct the 3D organizations of chromosomes. GEM-FISH fully exploits the complementary nature of FISH and Hi-C data constraints to improve the modeling process and reveal the finer details of the chromosome packing. In particular, it first uses both Hi-C and FISH data to calculate a TAD-level-resolution 3D model of a chromosome and reconstruct the 3D conformations of individual TADs using the intra-TAD interaction frequencies from Hi-C maps and the radii of gyration derived from FISH data. After that, an assembly algorithm is used to integrate the intra-TAD conformations with the TAD-level-resolution model to derive the final 3D model of the chromosome. We have demonstrated that GEM-FISH can obtain better 3D models than using Hi-C data only, with more accurate spatial organizations of TADs and compartments in the 3D space. In addition, we have shown that the final 3D models reconstructed by GEM-FISH can also capture the spatial proximity of loop loci and the colocalization of loci belonging to the same subcompartments. Based on our modeled 3D organizations of chromosomes, we have also found interesting patterns of the spatial distributions of super-enhancers on the three autosomes investigated (i.e., Chrs 20, 21, and 22). This novel finding can provide useful mechanistic insights into understanding the regulatory roles of super-enhancers in controlling gene activities.

## 2 Results

### 2.1 Integrating Hi-C and FISH data for modeling the 3D structure of a chromosome

We propose a divide-and-conquer based method, called GEM-FISH, to determine the 3D spatial organization of a chromosome through systematically integrating both Hi-C and FISH data. Our framework consists of the following three main steps (Fig. 1). First, we determine the 3D spatial arrangements of individual TADs at TAD-level resolution using both Hi-C and FISH data. Second, we compute the 3D coordinates of genomic loci within individual TADs using a sufficient number of geometric constraints derived from Hi-C data. During the elucidation of both intra-TAD and inter-TAD structures in the previous two steps, we also consider the prior biophysical knowledge of 3D polymer models to compute biophysically feasible and structurally stable structure of a chromosome. In the last step, the modeling results from the previous two steps are assembled through a series of translation, rotation, and reflection operations on individual TAD conformations. Below, we will describe the details of individual steps.

**Figure 1.**
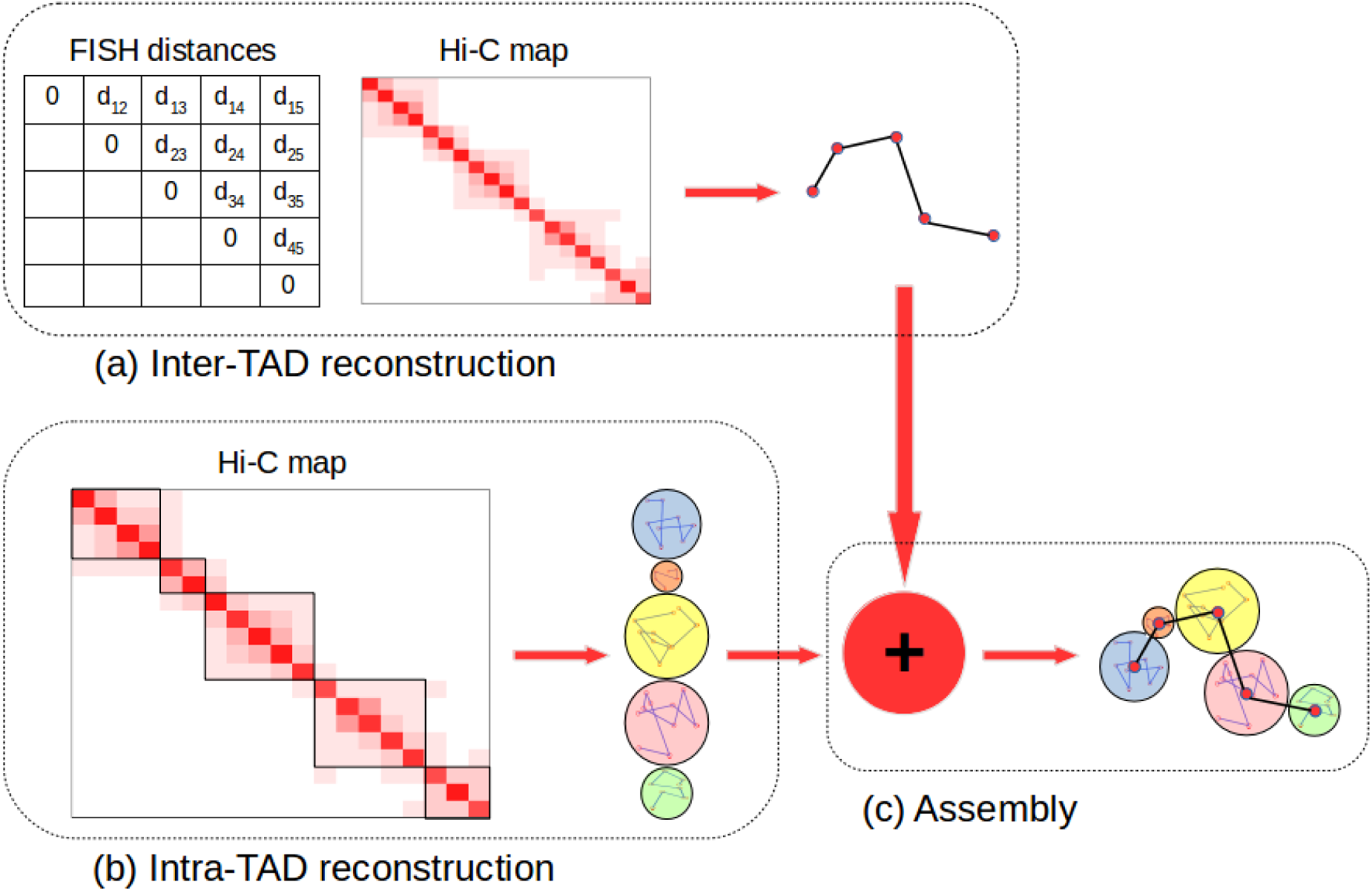
The schematic overview of GEM-FISH, which applies a divide-and-conquer strategy to reconstruct the 3D organization of a chromosome by systematically integrating both Hi-C and FISH data. (a) The 3D chromosome model at TAD-level-resolution is calculated by integrating Hi-C and FISH data as well as prior biophysical knowledge of a 3D polymer model. (b) The 3D conformations of individual TADs are determined using the intra-TAD geometric restraints derived from the input Hi-C map and prior biophysical knowledge of polymer models. (c) The final complete 3D structure of the chromosome is obtained by assembling the modeling results from the previous two steps, i.e., placing the previously determined intra-TAD conformations into the TAD-level-resolution model through translation, rotation, and reflection operations. More details can be found in the main text.

#### 2.1.1 Determining the 3D chromosome models at TAD-level resolution using both Hi-C and FISH data

We first calculate a relatively low-resolution (i.e., at TAD-level resolution) model of the chromosome of interest that is consistent with both input Hi-C and FISH data and also biophysically stable. Although a Hi-C map generally provides a relatively less number of geometric constraints between TADs than within individual TADs, these inter-TAD interaction frequencies can still provide useful restraints for pinning down the spatial arrangements of individual TADs at TAD-level resolution. On the other hand, FISH techniques directly image the 3D coordinates of different TADs from a number of cells, based on which we can also derive the average spatial distance between a pair of TADs over all cells. Overall, Hi-C and FISH data can provide useful and complementary restraints to determine the 3D organization of a chromosome at TAD-level resolution.

More specifically, to calculate the global chromosome structure at TAD-level resolution, we optimize a cost function that simultaneously incorporates the constraints derived from Hi-C data, FISH data, and prior biophysical knowledge about a 3D polymer model. In particular, the cost function *C_g_* is defined as,

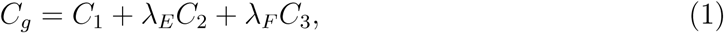
where *C*_1_, *C*_2_, and *C*_3_ stand for the cost terms corresponding to the restraints derived based on Hi-C data, prior biophysical knowledge, and FISH data, respectively, and *λ_E_* and *λ_F_* represent the corresponding coefficients that weigh the relative importance of individual terms.

The term *C*_1_ is defined using the same strategy as in [32], that is,

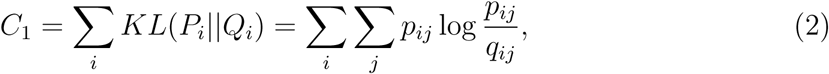
where *KL*(.) represents for the Kullback-Leibler (KL) divergence between two distributions, *p_ij_* represents the neighboring affinity between two genomic loci *l_i_* and *l_j_* in Hi-C space and *q_ij_* represents the probability that two 3D points *s_i_* and *s_j_* (corresponding to loci *l_i_* and *l_j_*, respectively) are close to each other in the reconstructed 3D chromosome model. Here, the neighboring affinity is derived according to the normalized interaction frequencies, that is,

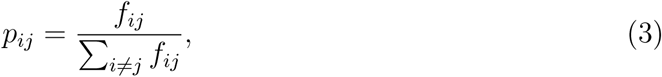
where *f_ij_* stands for the interaction frequency between the two genomic loci *l_i_* and *l_j_*. In addition, *q_ij_* is defined as follows,

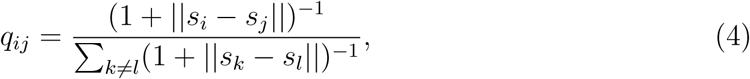
where ||.|| stands for the Euclidean distance between two 3D points.

In the cost function defined in Eq. 1, the term *C*_2_ represents the conformation energy of a 3D polymer model, which is defined using the same strategy as in [17,32]. The term *C*_3_ is a new term that we add to incorporate the average spatial distance constraints derived from FISH imaging data. Let *F_ij_* denote the average spatial distance between two TADs *t_i_* and *t_j_* measured from FISH experiments. Then *C*_3_ is defined as,

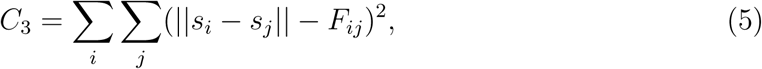
where *s_i_* and *s_j_* stand for the coordinates of the centers of TADs *t_i_* and *t_j_*, respectively.

More technical details about optimizing the cost function *C_g_* and selecting the optimal parameters *λ_E_* and *λ_F_* can be found in the “Methods” section.

#### 2.1.2 Determining the 3D chromosome conformations within individual TADs

Since FISH data only provide the geometric restraints on the 3D chromosome models at TAD-level resolution, and do not provide high-resolution information about the internal structure of each TAD, we cannot use them as pairwise distance restraints to determine the 3D coordinates of genomic loci within individual TADs. Nevertheless, we can still use FISH data to obtain a rough estimate of the radius of gyration (denoted by 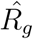) of every TAD (more details can be found in the “Methods” section). In principle, incorporating 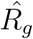 as an additional constraint to determine the 3D structures of intra-TAD chromosome fragments can further improve the modeling accuracy. Note that a similar scheme has also been used to incorporate FISH data to model the 3D genome structures from single-cell Hi-C data [38]. To incorporate the estimated radius of gyration into our modeling process, we define a new term *C*_4_,

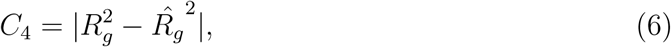
where *R_g_* stands for the radius of gyration of the reconstructed 3D model of the corresponding TAD and is calculated as follows,

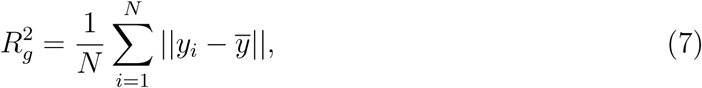
where *N* stands for the number of genomic loci in the 3D model, *y_i_* is the 3D coordinates of individual loci, 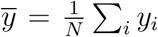, and ||.|| stands for the Euclidean distance between two loci in the reconstructed 3D model.

Then the cost function for calculating the local 3D chromosome structures within individual TADs is defined as,

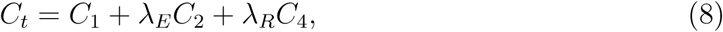
where *C*_1_ and *C*_2_ stand for the terms representing the constraints from Hi-C data and prior biophysical knowledge, respectively, and *λ_E_* and *λ_R_* stand for the coefficients that weigh the relative importance of the corresponding terms. More details about selecting the optimal parameters *λ_E_* and *λ_R_* can be found in the “Methods” section.

#### 2.1.3 Integrating the individual intra-TAD conformations with the global TAD-level-resolution model

The previously-computed global chromosome model at TAD-level resolution basically consists of a list of 3D points, each of which corresponds to the center of a TAD. We then translate the modeled internal conformations of individual TADs such that their centers match the corresponding points in the global structure.

In [6], it was observed that the Hi-C contact frequencies are inversely proportional to the 4*^th^* power of the corresponding mean spatial distances measured from FISH imaging data at TAD-level resolution. Here, we use the same strategy to estimate the spatial distance between two subsequent TADs from Hi-C data, i.e., between the last locus of a specific TAD and the first locus of the subsequent TAD (along the genome sequence). Through a series of further rotation and reflection operations on each TAD model, we minimize the difference between the spatial gap between two sequential TADs and its estimate from Hi-C data. More details about assembling the intra-TAD structures of a chromosome can be found in the “Methods” section.

### 2.2 GEM-FISH yields more accurate three-dimensional chromosome models

To evaluate the modeling performance of GEM-FISH, we used it to compute the 3D models of Chromosomes 20, 21, 22, and X of human diploid IMR90 cells for which both Hi-C [5] and FISH [6] data are available. For Chromosome X, we calculated both its active and inactive states 3D models, denoted by ChrXa and ChrXi, respectively. We assessed the accuracy of our reconstructed models by measuring the relative error (denoted by *RE*) in the distance between each pair of TADs, which is defined as,

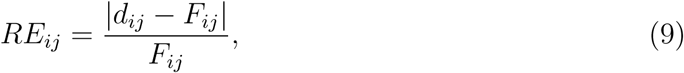
where the term *F_ij_* stands for the average distance between TADs *i* and *j* obtained from FISH imaging data, and the term *d_ij_* denotes the distance between the centers of two TADs *i* and *j* in case of the low-resolution (i.e., TAD-level-resolution) chromosome model or the average pairwise distance over all pairs of genomic loci between TADs *i* and *j* in case of the final complete model.

Fig. 2(a) and 2(b) show the TAD-level-resolution model and the corresponding final model of an example chromosome (i.e., Chr21). We calculated the relative errors of both TAD-level-resolution and final models resulting from GEM-FISH using both Hi-C and FISH data (Fig. 2(c) and 2(e)). For comparison, we also calculated the relative errors of both TAD-level-resolution and final models reconstructed by GEM [32] using only Hi-C data (Fig. 2(d) and 2(f)). We found that the integration of FISH constraints with Hi-C data significantly decreased the relative errors in the spatial distances especially between TADs far away along the genome that usually have relatively low Hi-C contact frequencies (Fig. 2(c) and 2(d)). In addition, when compared to the TAD-level-resolution models, the relative errors in the spatial distances between adjacent TADs along the genome slightly decreased in the final model (see the diagonal elements in Fig. 2(e) and 2(f)), which basically indicated that our method was able to compute the correct relative orientations of adjacent TADs.

**Figure 2.**
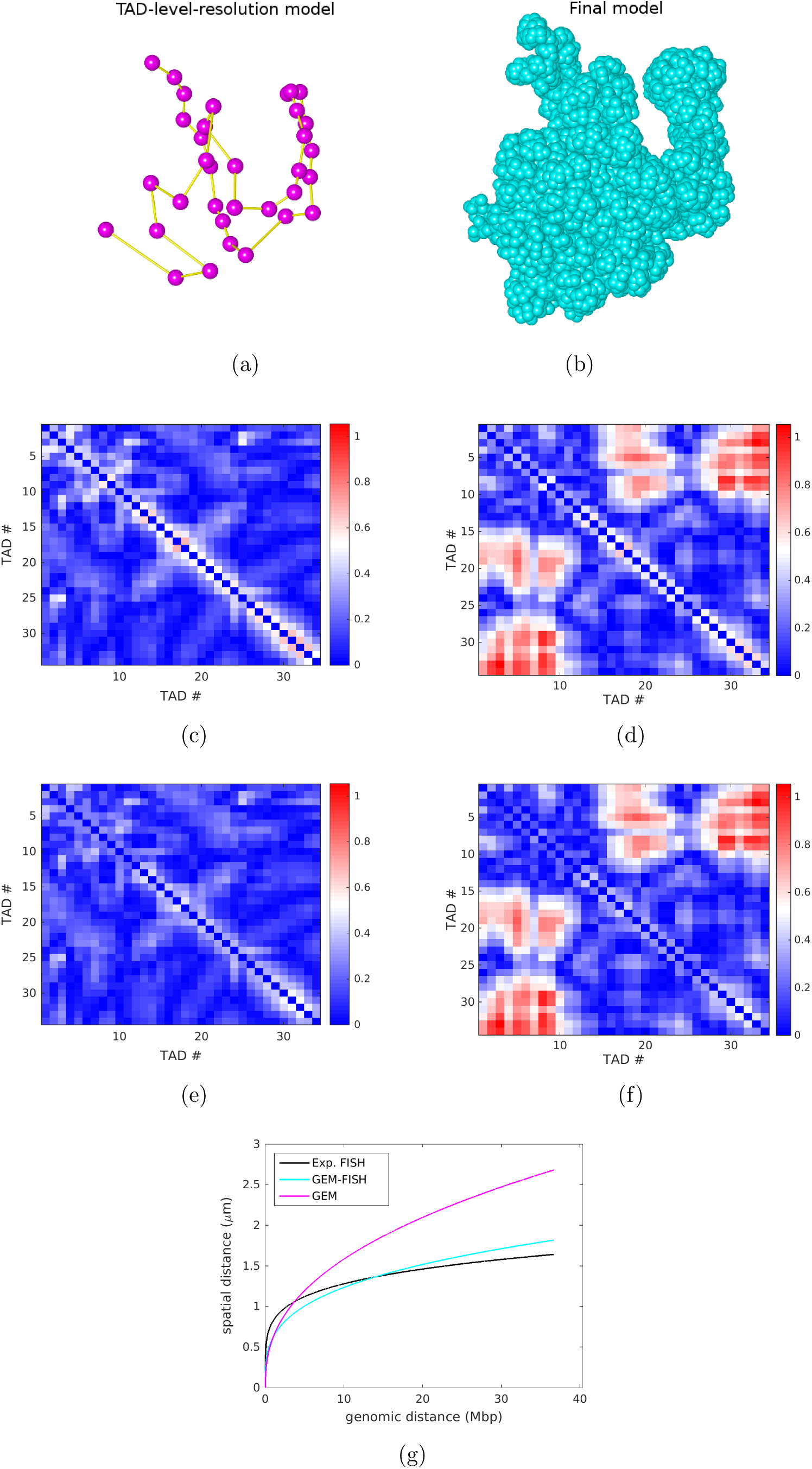
The modeling results of human Chromosome 21 (Chr21). (a) The TAD-level-resolution 3D structure of Chr21 calculated by GEM-FISH, where each dot represents the center of a TAD. (b) The final 3D structure of Chr21 reconstructed by GEM-FISH. The visualization in (a) and (b) was performed using UCSF Chimera [48]. (c and d) The relative error matrices of the TAD-level-resolution models computed by GEM-FISH using both Hi-C and FISH data, and by GEM using only Hi-C data, respectively. (e and f) The relative error matrices of the final models computed by GEM-FISH using both Hi-C and FISH data, and GEM using only Hi-C data, respectively. (g) The curves of spatial vs. genomic distances between TADs, which were derived from the experimental FISH data, and the final 3D models reconstructed by GEM-FISH (using both Hi-C and FISH data), and by GEM (using only Hi-C data), respectively.

We also compared the curves of the spatial vs. genomic distances between TADs for the models reconstructed by GEM-FISH using both Hi-C and FISH data, and GEM [32] using only Hi-C data, respectively, to those derived directly from the experimental FISH data [6]. We found that the relationships between genomic vs. inter-TAD spatial distances in the final 3D models computed by GEM-FISH were much closer to those derived from the FISH experimental data than the 3D structures derived from GEM using the Hi-C data alone, especially for the regions with relatively large genomic distances (Fig. 2(g)). Table 1 summarizes the relative errors obtained for all five tested chromosomes, and Figures S1-S4 show the corresponding modeling results for Chrs 20, 22, Xa, and Xi, respectively. All these results indicated that incorporating FISH data can significantly improve the accuracy of modeling the 3D organizations of chromosomes.

**Table 1.** The average relative errors of the TAD-level-resolution and final models reconstructed by GEM-FISH using both Hi-C and FISH data, and by GEM using Hi-C data only for Chrs 20, 21, 22, Xa, and Xi.

|  | TAD-level-resolution model<br>by GEM-FISH | Final model<br>by GEM-FISH | TAD-level-resolution model<br>by GEM | Final model<br>by GEM |
| --- | --- | --- | --- | --- |
| Chr20 | 0.18 | 0.16 | 0.31 | 0.30 |
| Chr21 | 0.16 | 0.14 | 0.31 | 0.30 |
| Chr22 | 0.17 | 0.16 | 0.22 | 0.20 |
| ChrXa | 0.17 | 0.16 | 1.36 | 1.37 |
| ChrXi | 0.22 | 0.21 | 1.92 | 1.93 |

### 2.3 The 3D chromosome models reconstructed by GEM-FISH have accurate compartment partition

It has been widely observed that chromosomes are partitioned into two compartments (i.e., A and B) based on Hi-C maps [2]. TADs within the same compartment are generally spatially closer to each other and have higher contact frequencies than those belonging to different compartments. Using FISH experiments, Wang et al. [6] observed that the two compartments are typically spatially arranged in a polarized fashion relative to each other. In addition, they showed that individual compartments are relatively enriched with different epigenetic marks.

Following the same strategy as in [6], we assigned TADs to A/B compartments for the 3D chromosome models calculated by GEM-FISH (using both Hi-C and FISH data) and GEM [32] (using only Hi-C data). For the examined autosomes (i.e., Chrs 20, 21, and 22), the average accuracy of assigning TADs of the 3D models reconstructed by GEM-FISH to A/B compartments was 89.6% vs. 81.0% for those models calculated by GEM (Table 2, Fig. 3). We found that the 3D models computed by GEM-FISH displayed approximately similar relative enrichment patterns of different epigenetic marks in A/B compartments for the three autosomes, which were close to those derived from experimental FISH data (Fig. 4). This observation indicated that the few TADs that were wrongly assigned to the A and B compartments in the models reconstructed by GEM-FISH probably had more noisy epigenetic properties of one compartment over the other. On the other hand, the difference was relatively more obvious for the 3D model of Chr20 calculated by GEM (Fig. 4(a)). This was likely due to the relatively low accuracy in assigning TADs to A/B compartments on this model when using only Hi-C data to reconstruct its 3D chromosome structure (73.3%, Table 2).

**Table 2.** The accuracy of assigning TADs of the 3D chromosome models calculated by GEM-FISH (which uses both Hi-C and FISH data) and GEM (which uses only Hi-C data) to the A and B compartments. The table entries are filled with the format of “number of correctly assigned TADs/total number of TADs”.

|  | GEM-FISH | GEM |
| --- | --- | --- |
| Chr20 | 28/30 | 22/30 |
| Chr21 | 32/34 | 30/34 |
| Chr22 | 22/27 | 22/27 |
| ChrXa | 39/40 | 35/40 |
| ChrXi | 37/40 | 24/40 |

**Figure 3.**
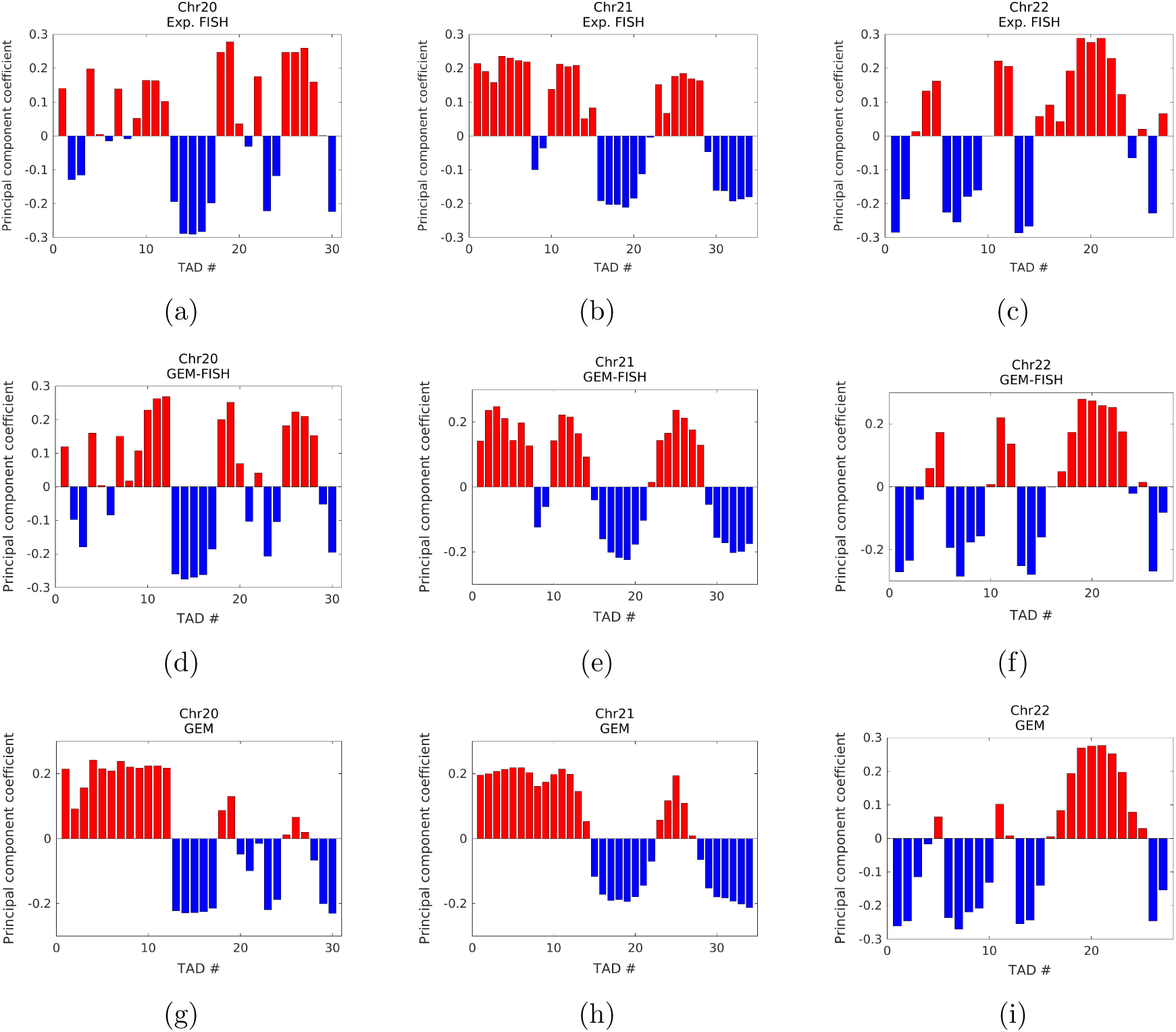
Assignment of TADs to A/B compartments for Chrs 20, 21, and 22. (a, b, and c) Assignment of TADs for Chrs 20 (a), 21 (b), and 22 (c) using the experimental FISH data (obtained from [6]). (d, e, and f) Assignment of TADs of the 3D chromosome models calculated by GEM-FISH using both Hi-C and FISH data for Chrs 20 (d), 21 (e), and 22 (f). (g, h, and i) Assignment of TADs of the 3D chromosome models calculated by GEM using only Hi-C data for Chrs 20 (g), 21 (h), and 22 (i).

**Figure 4.**
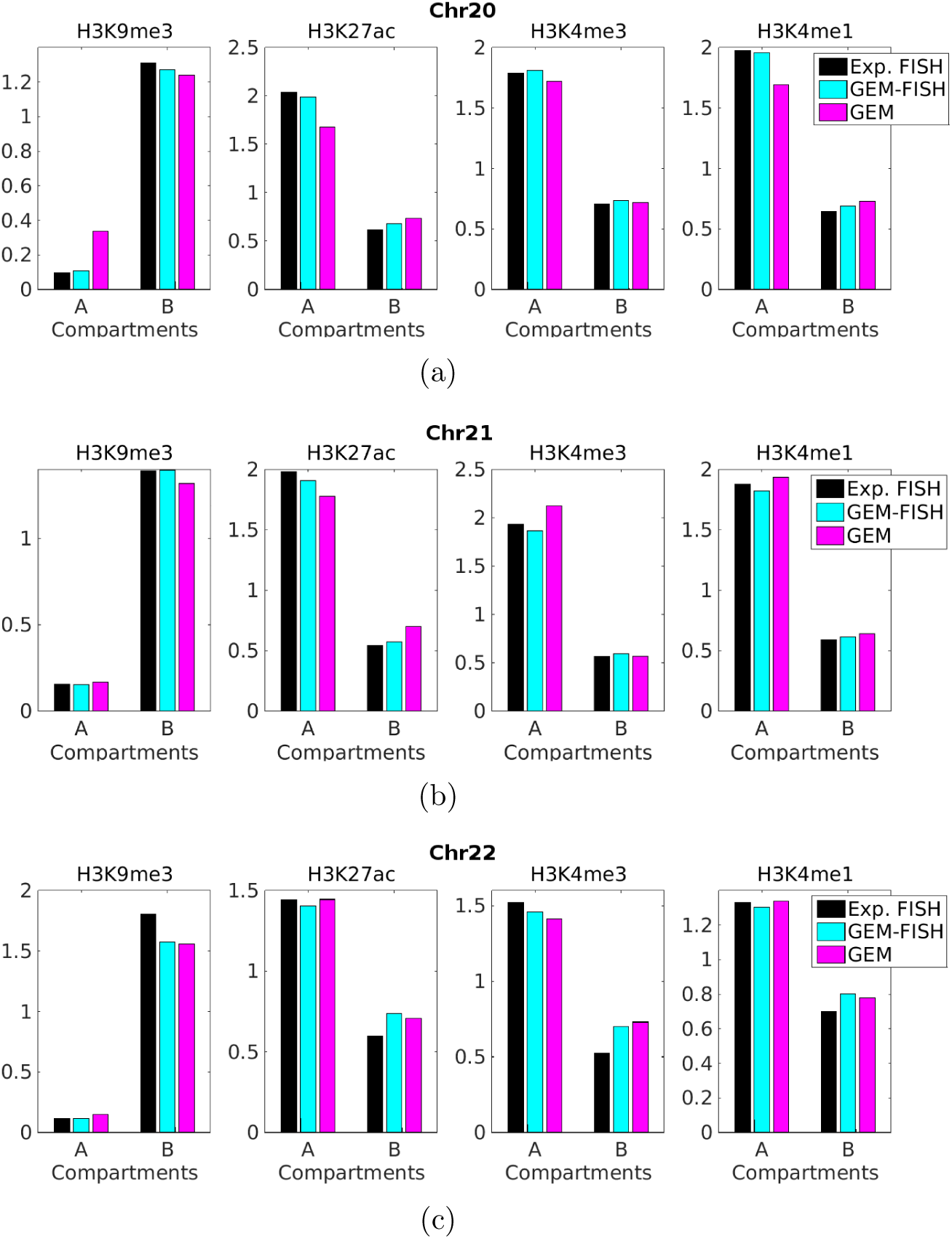
Relative enrichment of different histone marks, including H3K9me3, H3K27ac, H3K4me3, and H3K4me1, on compartments A and B of Chr20 (a), Chr21 (b), and Chr22 (c) for different schemes of assigning TADs to A/B compartments, including using the experimental FISH data and the 3D chromosome models reconstructed by GEM-FISH (using both Hi-C and FISH data) and GEM (using only Hi-C data).

The final 3D models calculated by GEM-FISH for ChrXa and ChrXi were clearly different and can be easily distinguished through visual inspection (Fig. 5(a) and 5(b)). The 3D model of ChrXi was notably more compact compared to that of ChrXa, which was consistent with its inactive nature. The quantitative comparison of the compactness of the 3D models of ChrXi and ChrXa showed that the densities of the TADs of ChrXi were significantly higher than those of ChrXa (Fig. 5(c)).

**Figure 5.**
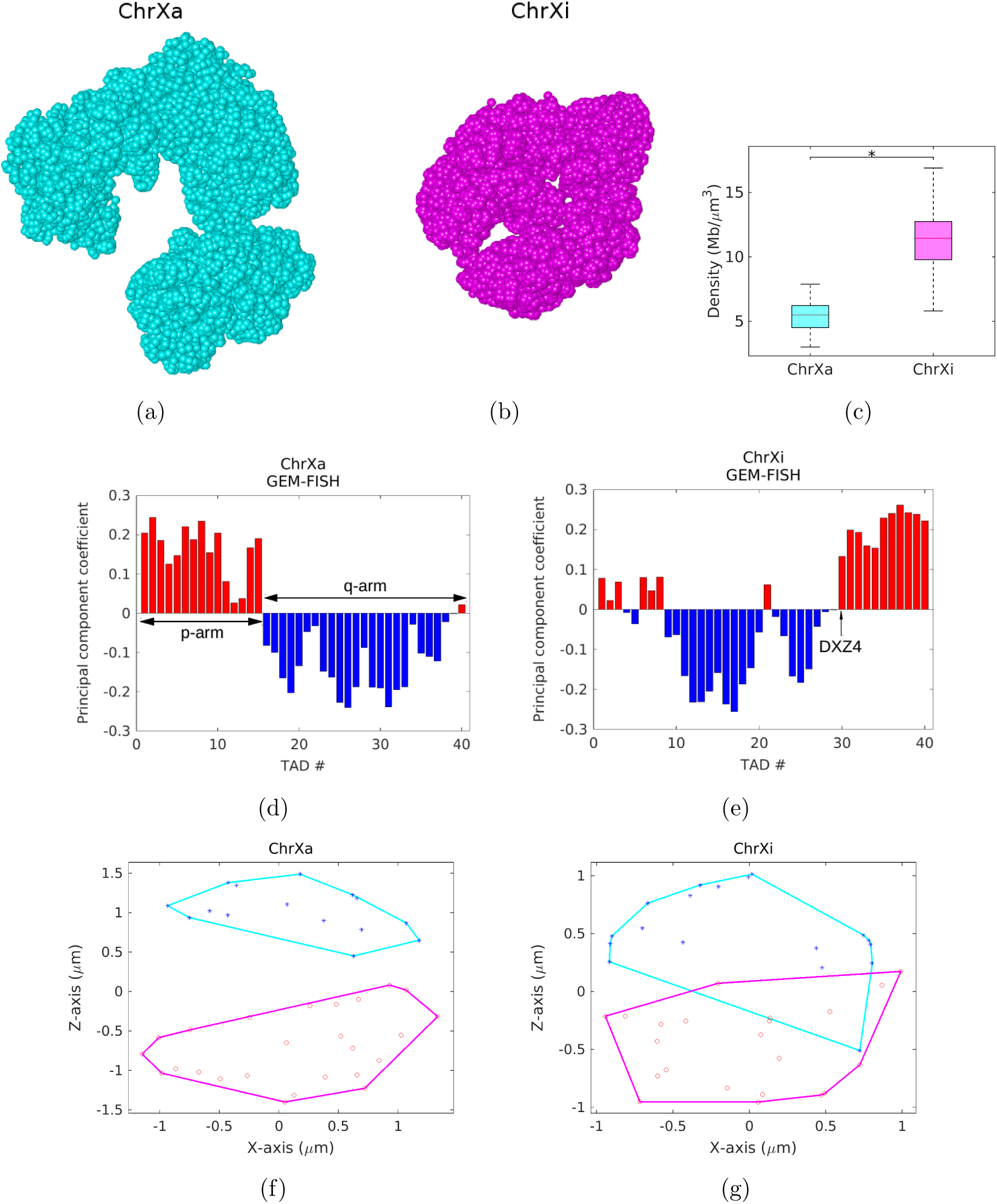
The modeling results of GEM-FISH for the human X-Chromosome (including both active state ChrXa and inactive state ChrXi). (a and b) Visualization of the final 3D models of ChrXa (a) and ChrXi (b) in UCSF Chimera [48]. (c) Comparison of the compactness of TADs between the 3D models of active (ChrXa) and inactive (ChrXi) states of human X-Chromosome. *: *p*-value < 10^−13^, one-tailed Wilcoxon rank-sum test. (d and e) The assignment of TADs of ChrXa (d) and ChrXi (e) to the A/B compartments. (f and g) Projection of the 3D convex hull plots of the A and B compartments of ChrXa (f) and ChrXi (g) to the XZ plane.

Although Wang et al. [6] observed that the X-Chromosome can be partitioned into two compartments, they found that the compartmentalization scheme was different for its active state ChrXa and inactive state ChrXi. The two compartments of ChrXa corresponded to its p and q arms. For ChrXi, there were two continuous compartments separated along the genomic sequence by the DXZ4 macrosatellite. Here, the 3D models calculated by GEM-FISH (Fig. 5(a)) resulted in 97.5% and 92.5% accuracy in assigning TADs to the two compartments for ChrXa and ChrXi, respectively (Table 2). More importantly, in the 3D models derived from GEM-FISH, the separation position between the two compartments was correctly captured for both ChrXa and ChrXi (Fig. 5(d) and 5(e)). On the other hand, since so far only the non-allele-specific Hi-C maps (which do not distinguish between ChrXa and ChrXi) were available for the IMR90 cell line [5] (to our best knowledge), the 3D models calculated by GEM (which only takes Hi-C data as input) resulted in lower accuracy in assigning their TADs to the two compartments (87.5% and 60.0% for ChrXa and ChrXi, respectively). The significant improvement in the accuracy of compartmentalization after integrating both Hi-C and FISH data into the modeling process especially for the X-Chromosomes showed that the FISH data can play an important role in deriving more accurate spatial organizations of TADs for individual chromosomes.

As in [6], we also examined how the two compartments of the X-Chromosomes were arranged spatially relative to each other. Our modeling results from GEM-FISH showed that the two compartments of all the obtained 3D chromosome models were placed in a polarized fashion relative to each other, with the degree of polarization in ChrXi less than that in ChrXa, which was consistent with the previous finding in [6] (Fig. 5(f), 5(g), and S5). In addition, we found that the pairwise spatial distances between TADs within the same compartment were significantly smaller than those between TADs across different compartments for all five tested chromosomes (Fig. 6(a)), which was also consistent with the previous result [2].

**Figure 6.**
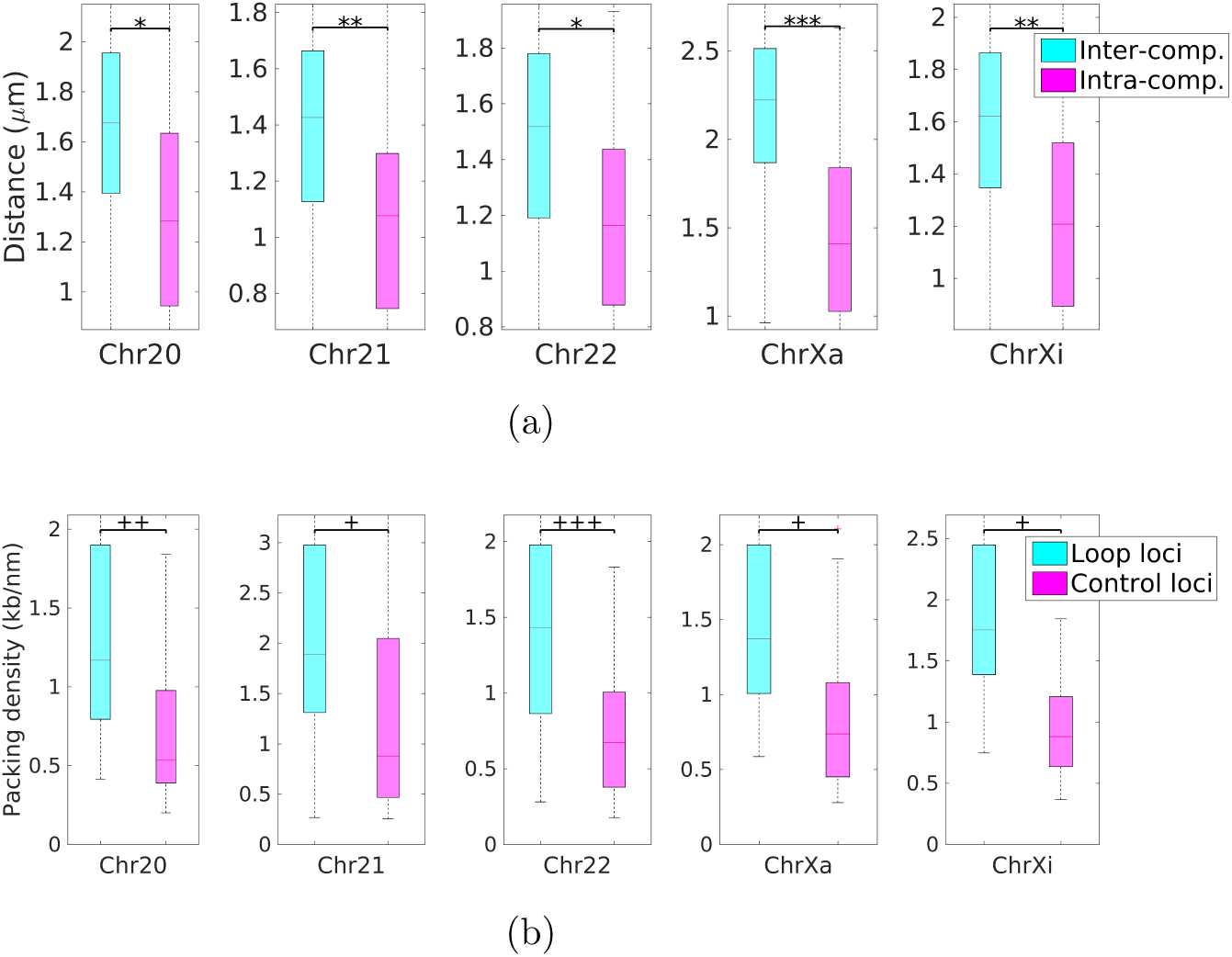
Comparisons of the pairwise TAD distances from the same compartment vs. from the different compartments, and the packing densities between loop loci vs. between control loci. (a) Comparisons of the pairwise distances between TADs within the same compartment vs. those between TADs from different compartments. *: *p*-value < 10^−26^, **: *p*-value < 10^−46^, ***: *p*-value < 10^−122^. (b) Comparisons of the packing densities between loop loci vs. between control loci. +: *p*-value < 10^−4^, ++: *p*-value < 10^−8^, +++: *p*-value < 10^−11^. All tests were performed using the one-tailed Wilcoxon rank-sum test.

### 2.4 Validating the loop conformations derived from GEM-FISH

Using their developed program HiCCUPS, Rao et al. annotated the loops in the genomes of nine different cell types from the high-resolution (1~10 Kbps) Hi-C maps [5]. To validate these high-resolution chromatin loops of the 3D models derived from GEM-FISH, we first used the lists of the loops of Chr20, Chr21, Chr22, and ChrX of the IMR90 cell line annotated from [5] to obtain the genomic positions of the corresponding loop loci. We then calculated the spatial distance between every two loop loci in our final 3D model computed by GEM-FISH. For each annotated loop, we also computed the spatial distance between one of the two loop anchor loci and a control locus that was located at the same genomic distance but on the opposite side of the genome. We found that the DNA packing densities of the regions between loop anchor loci were significantly higher than those of the regions between control loci for all five tested chromosomes (Fig. 6(b)). Such result provides further support to the final 3D models reconstructed by GEM-FISH.

### 2.5 Analyses of subcompartments

It has been observed that there are at least six nuclear subcompartments defined based on their long-range interaction patterns [5]. Two subcompartments associate with compartment A, hence called A1 and A2, and the other four subcompartments associate with compartment B, hence called B1, B2, B3, and B4. The genomic loci belonging to different subcompartments tend to exhibit distinct genomic and epigenomic properties. For instance, those loci belonging to subcompartments A1 and A2 are enriched with activating chromatin marks, such as H3K27ac, H3K36me3, H3K4me1, and H3K79me2 [5]. On the other hand, the loci belonging to subcompartment B1 correlate positively with the repressive mark H3K27me3 and negatively with the activating mark H3K36me3, while the loci belonging to subcompartment B2 lack all the marks mentioned above [5].

Rao et al. [5] mainly used the interchromosomal interaction frequency information from Hi-C data to annotate the subcompartments. Unfortunately, due to the sparsity of the interchromosomal Hi-C maps for most of the cell lines inspected, they only annotated the subcompartments of the cell line GM12878. Recently, Di Pierro et al. [39] used the chromatin immunoprecipitation-sequencing (ChiP-Seq) data of histone marks and other protein-binding experiments to infer the subcompartment types of genomic loci. They partitioned the chromosomes into individual bins of size 50 Kbps and assigned each bin with a discrete score from 1 to 20 representing the enrichment of a certain chromatin mark. They then used a neural network that takes the scores of different chromatin marks of a certain bin as inputs to infer its subcompartment type. In our work, we followed the same strategy of assigning each 50 Kbps bin with a discrete score representing the enrichment of a chromatin mark. Here, we mainly considered the subcompartment types B1 and B2, particularly because they are expected to have different folding properties due to their distinct epigenomic content. For instance, the genomic regions belonging to subcompartment B1 with a repressive nature are expected to be more densely packed than those belonging to subcompartment B2 with an inactive nature [34]. Instead of building a neural network for classification as in [39], we used a simple method to determine the subcompartment type of a specific genomic region. In particular, for the three autosomes Chr20, Chr21, and Chr22, we considered the loci that have a score less than three for the aforementioned chromatin marks (i.e., H3K27me3, H3K36me3, H3K27ac, H3K4me1, and H3K79me2) to be assigned to subcompartment B2. On the other hand, we considered those loci that have a score more than or equal to ten for H3K27me3 and less than three for H3K36me3 to be assigned to subcompartment B1. The threshold three was chosen to filter experimental noise that may affect our analysis, and the threshold ten was used to choose the enrichment scores above the median. We found that the genomic loci belonging to a certain subcompartment tend to colocalize in the 3D models reconstructed by GEM-FISH (Fig. 7(a), 7(c), and 7(e)), which agrees well with the previous finding in [5]. We also calculated the densities of regions belonging to the two subcompartments in the reconstructed 3D models and found that the densities of the regions from subcompartment B1 were significantly higher than those of the regions from subcompartment B2 (Fig. 7(b), 7(d), and 7(f)). Such a finding was also consistent with the previous result in [34].

**Figure 7.**
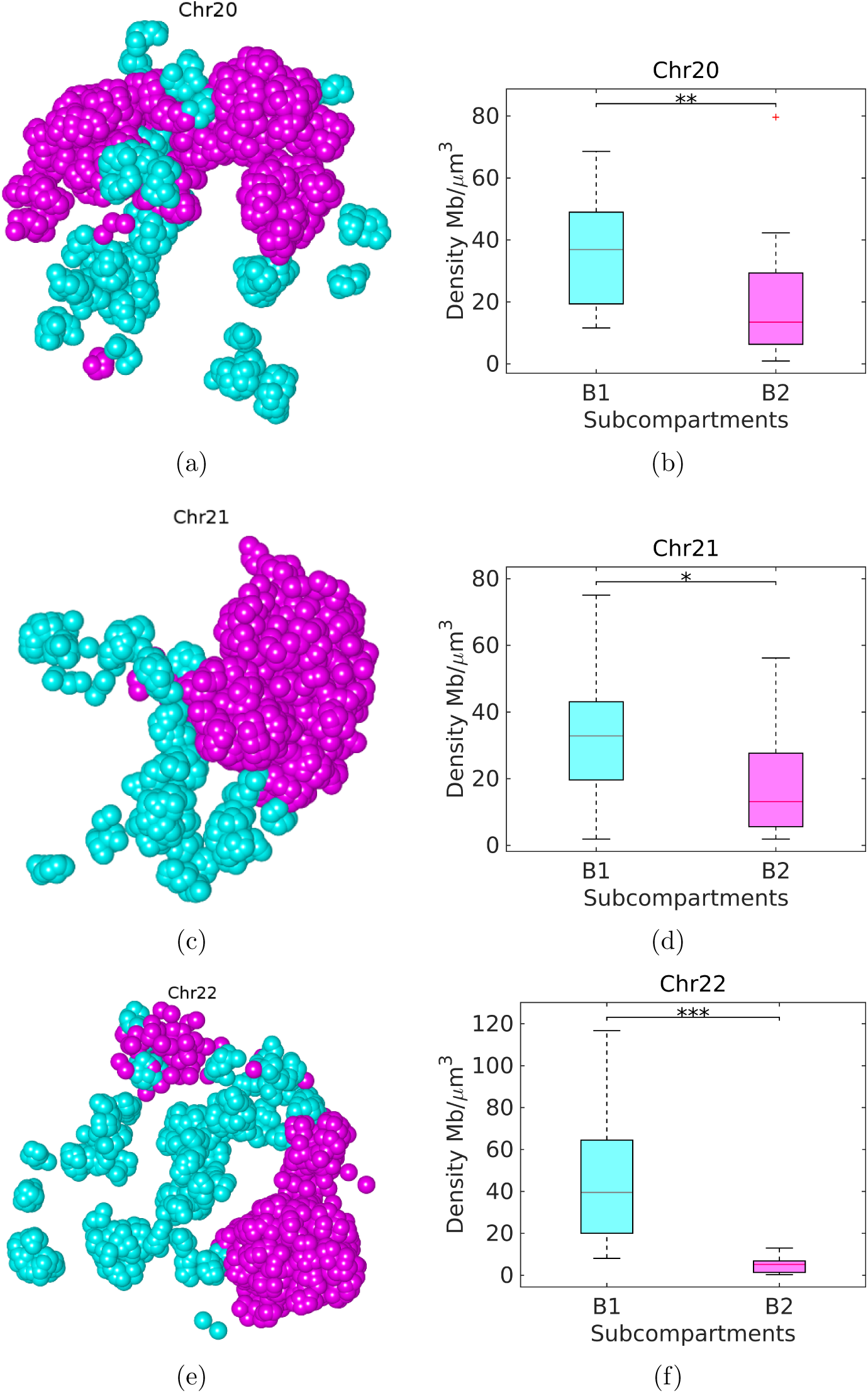
Regions belonging to the same subcompartment tend to colocalize in the 3D space. (a, c, and e) Visualization of the regions belonging to subcompartments B1 (cyan) and B2 (magenta) in the 5 Kbps-resolution 3D models reconstructed by GEM-FISH for Chr20, Chr21, and Chr22, respectively. Only regions that belong to either B1 or B2 are shown. The visualization was performed using UCSF Chimera [48]. (b, d, and f) Boxplots on the densities of regions belonging to subcompartments B1 and B2 for Chr20, Chr21, and Chr22, respectively. *: p-value < 0.007, **: p-value < 10^−4^, ***: p-value < 10^−5^. All tests were performed using the one-tailed Wilcoxon rank-sum test.

### 2.6 The novel patterns of the spatial distributions of super-enhancers

Super-enhancers are a group of enhancers that are in close genomic proximity and span a genomic interval in a range of tens of kilobase pairs. A key feature that distinguishes super-enhancers from common enhancer is the relatively high enrichment of specific transcription coactivators such as mediator Med1 or activating histone marks such as H3K27ac. They are usually found close to the cell-type-specific genes that define the cell identity and regulate their expression [40,41].

We first obtained the positions of super-enhancers of Chrs 20, 21, and 22 from the super-enhancer database dbSUPER [40], and then examined their locations in the final 3D models calculated by GEM-FISH. We found that super-enhancers tend to lie on the surfaces of the reconstructed 3D chromosome models (Fig. 8), which was consistent with the previous finding that active regions usually tend to lie on the periphery of the chromosome territory [39,42]. In addition, for Chr21, we found that four of its five super-enhancers lie in the G-band q22.3. After closely examining this band, we found that it covers about 40% of the currently known protein-coding genes in Chr21 [43], although it forms only less than 12% of the size of the chromosome. From the visualization of this band in the final 3D model reconstructed by GEM-FISH, we found that it formed an arm-like structure extending out from the chromosome body, and appeared to be accessible from all directions (Fig. 8(b)), which was consistent with the active nature of the whole band. By contrast, the discrimination of this gene-rich band was not that obvious in the final model reconstructed by GEM using Hi-C data alone (Fig. S6), in which other parts of the chromosome also formed arm-like structures accessible from all directions. These novel patterns of the spatial distributions of super-enhancers can not only provide important insights into revealing their functional roles in gene regulation, but also support the hypothesis that local interactions between genomic loci are the driving forces that lead to particular chromosome conformations [42,44,45].

**Figure 8.**
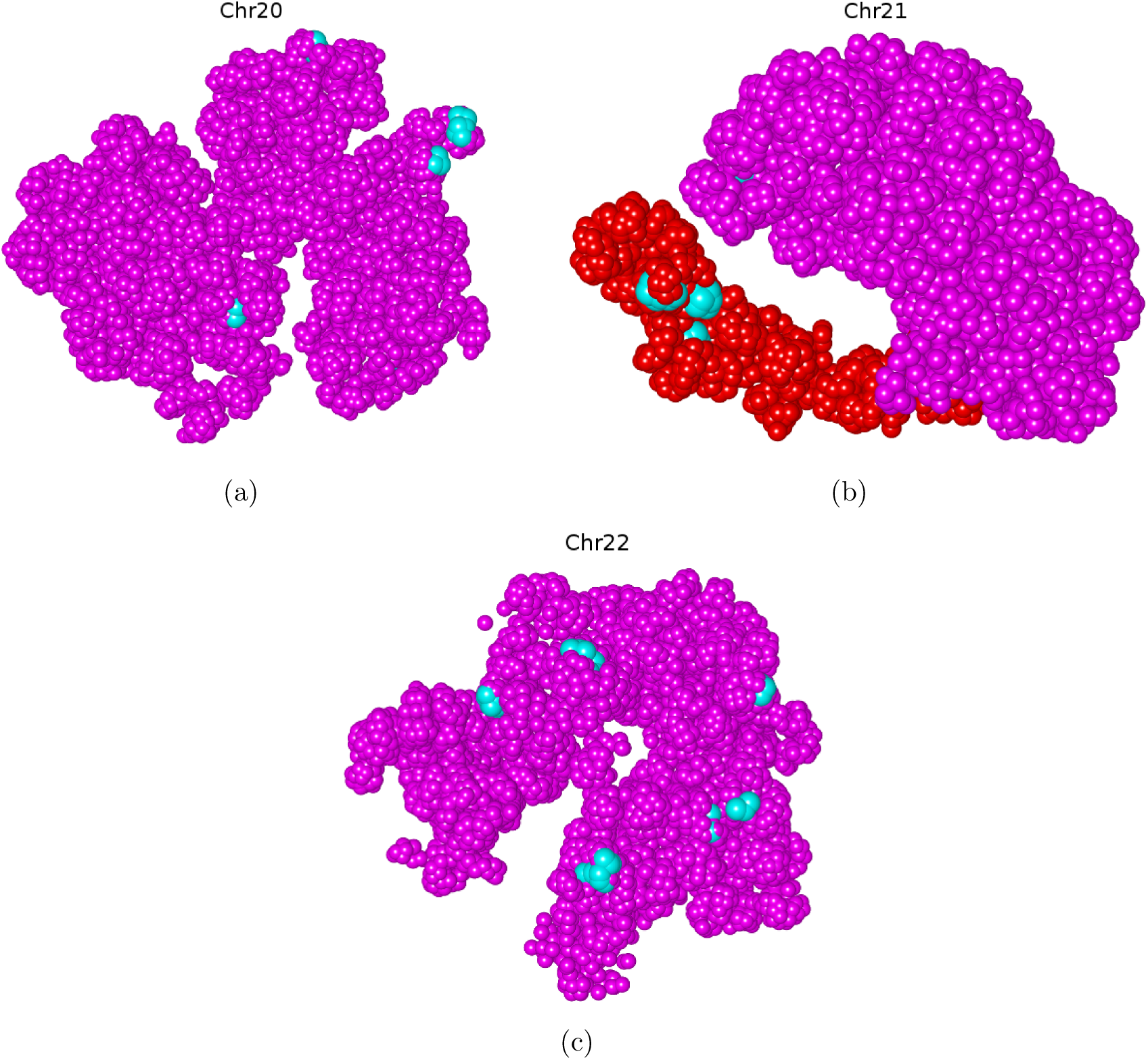
Super-enhancers tend to lie on the surface of the 3D models of chromosomes. (a-c) Super-enhancers (cyan) lie on the surfaces of the 3D models reconstructed by GEM-FISH for Chrs 20, 21, and 22, respectively. For Chr21 (b), four of the five super-enhancers were found in the gene-rich G-band q22.3 region, which is shown in red. The visualization was performed using UCSF Chimera [48].

## 3 Discussion and Conclusions

Both Hi-C and FISH techniques have been widely used to study the 3D genome structure. Hi-C data provide the contact frequencies between genomic loci and can be interpreted as a measure of how frequently a pair of two genomic loci come close to each other in the 3D spatial space, while FISH directly measures the spatial distances between genomic loci through the imaging techniques. Usually, both Hi-C and FISH provide consistent measures, i.e., the Hi-C contact frequency between a pair of genomic loci is normally inversely proportional to their spatial distance measured by FISH. In some cases, however, the results from these two methods may seem to contradict each other. This apparent contradiction has brought up an interesting research problem, driving scientists to read the results of these two techniques in an attempt to reconcile them [37,46].

Due to the cell-to-cell variability, the spatial distance between two genomic loci can change from one cell to another. The results from FISH experiments can be used to discover the distribution of the spatial distance between a pair of genomic loci. On the other hand, the Hi-C experiments capture only the interaction between two genomic loci if they are close to each other in the 3D space, which may only happen in a small population of the cells inspected. In [37], Fudenberg and Imakaev examined the Hi-C and FISH data obtained from [5] and showed that, although the median spatial distance of each loop-control pairs of loci changes concordantly with the corresponding Hi-C signal, this concordance does not always hold when comparing loop loci and control loci (i.e., loci that do not form loops) from different pairs. However, they noticed that the median spatial distance changes concordantly with the Hi-C signals for all pairs of control loci.

In the FISH data [6] used in our tests, the imaged loci span the central 100 Kbps segments of individual TADs. Each pair of these imaged loci belongs to different TADs and unlikely form a loop. Thus, the pairs of imaged loci were basically equivalent to those control loci investigated in [37]. To examine the consistency between Hi-C and FISH data used in our tests, we measured how often the median distances between TAD pairs measured from FISH varied concordantly with the Hi-C contact frequencies. As shown in Table S1, the average consistency between Hi-C and FISH data for Chr20, Chr21, and Chr22 was 82.03%, much higher than that of the control maps. Thus, we can conclude that both Hi-C and FISH data used in our study were reasonably consistent with each other.

In this study, we presented a new divide-and-conquer based method for modeling the 3D organizations of chromosomes. Our approach integrates both Hi-C and FISH data, as well as our current biophysical knowledge about a 3D polymer model. These different sources of information provide complementary constraints that allow the reconstruction of more accurate 3D models that can capture both global and local geometric features of chromosomes. On the one hand, the global features were validated through the highly accurate assignment of TADs to A/B compartments, the reasonable placement of compartments in a polarized fashion relative to each other, and the clear proximity of TADs within the same compartment. On the other hand, the local features were validated through the proximity of loop loci, the colocalization and different epigenomic properties of the genomic segments belonging to the same subcompartment. In addition, the 3D models of chromosome reconstructed by our method revealed interesting patterns of the spatial distributions of super-enhancers. Such a novel finding will provide important hints for further investigating the functional roles of super-enhancers in controlling gene activities.

In general, every source of available data has its own merits and limitations. Integrating multiple sources of data constraints can help fully exploit their benefits and overcome their weaknesses during the 3D chromosome structure modeling process. On the way to calculate the whole 3D genome model, more data sources will be needed to further increase the accuracy of the reconstructed structure, and also compensate the limited availability and modeling power of existing input data. For instance, the geometric constraints derived from lamina-DamID experiments can also be used to infer the proximity of a chromatin region to the nuclear envelope [47]. In addition, the epigenomic profiles derived from ChiP-Seq can provide additional useful information to reconstruct the 3D architectures of chromosomes [39]. In principle, our framework can be easily extended to integrate all these different types of data constraints for modeling the 3D structures of chromosomes, which will thus further improve our current understanding of the underlying functional roles of 3D genome folding in gene regulation.

## 4 Methods

### 4.1 Integrating FISH data into the modeling of individual TAD structures

The FISH data of the IMR90 cell line obtained from [6] did not provide any direct information about the pairwise distances between genomic loci inside individual TADs. Nonetheless, we can still derive rough estimates of the volume, size, and radius of gyration *R_g_* of every TAD using the available FISH data, which can still provide useful information for modeling the 3D structures of individual TADs.

The FISH data derived from [6] of a specific chromosome, denoted by *Chr_i_*, include a set of *n* structures, where *n* stands for the number of cells in which *Chr_i_* was imaged. Each structure of *Chr_i_* consists of *m* three-dimensional points, each corresponding to a TAD probed in that chromosome. The genomic regions in *Chr_i_* probed by FISH experiments in [6] either cover all its TADs (as in Chr21 and Chr22) or span approximately uniform intervals in the chromosome (as in Chr20 and ChrX).

For each of the *n* structures of *Chr_i_*, based on the FISH data measured in [6], we first derive an estimate of its volume (i.e., the volume of the minimum bounding box (mbb) circumscribing that structure) and its size (i.e., the length of the longest side of the mbb), and also calculate its radius of gyration, denoted by *R_g_*. We then plot histograms to show the distributions of volumes and sizes for individual structures of *Chr_i_* and select the values corresponding to the highest values to be the expected volume and size of that chromosome. We also infer an approximate linear relation between the size of the chromosome and its radius of gyration (Fig. S7 - S11).

We use 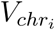 and 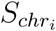 to denote the expected volume and size of *Chr_i_*, respectively. Given that the volume of a genomic region basically scales linearly with its genomic length [10], a rough estimate of the volume of *TAD_j_* in *Chr_i_* can be derived by:

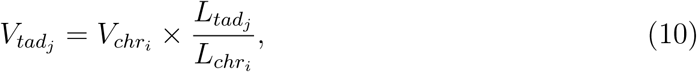
where 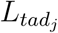 and 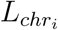 stand for the genomic lengths of *TAD_j_* and *Chr_i_*, respectively.

Since the ratio between the sizes of *TAD_j_* and *Chr_i_* (which are denoted by 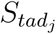 and 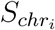, respectively) is approximately equal to the third power of the corresponding ratio of their volumes, we have

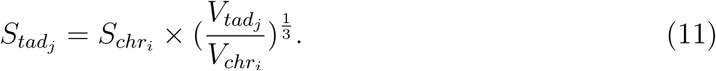

Using the approximate linear relation that we previously inferred between the size of the chromosome and its radius of gyration (Fig. S7 - S11), we can obtain an estimate of the radius of gyration 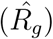 of *TAD_j_*. After that, we can place this estimated value into Eq. 6 and use it as an additional constraint to model the 3D structure of *TAD_j_*.

### 4.2 Optimization of the cost function

We use gradient descent to minimize the cost functions *C_g_* and *C_t_* defined in Eqs. 1 and 8, respectively. The gradient of *C_g_* with respect to the coordinate *s_i_* is calculated as follows,

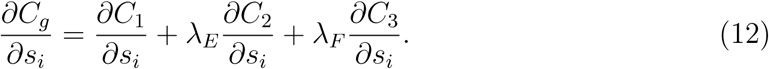

More details about the calculation of 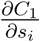 and 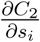 can be found in [32]. In addition, 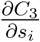 is calculated as follows,

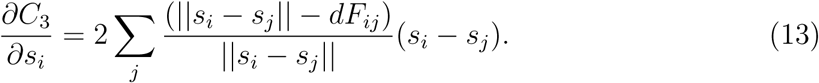

Similarly, the gradient of *C_t_* with respect to the coordinate *y_i_* is calculated as follows,

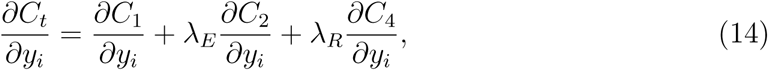

where 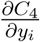 is calculated as follows,

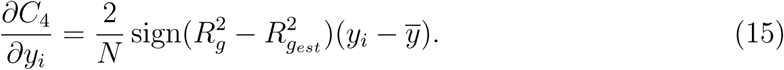

### 4.3 Parameter selection

In GEM-FISH, we optimize the cost function *C_g_* in Eq. 1 to calculate the TAD-level-resolution 3D model of the whole chromosome, and the cost function *C_t_* in Eq. 8 to calculate the models of individual TADs. Each of these two cost functions has two parameters (*λ_Ε_* and *λ_F_* in Eq. 1, and *λ_Ε_* and *λ_R_* in Eq. 8). These parameters need to be chosen in a principled way that the calculated models best interpret the input Hi-C and FISH data, and the prior knowledge of a 3D polymer model. From the available FISH data, we can obtain rough estimates of the volumes of the whole chromosome and individual TADs (see Section 4.1). Following the same strategy as in [32], we can select a pair of parameter values that maximize the following scoring function,

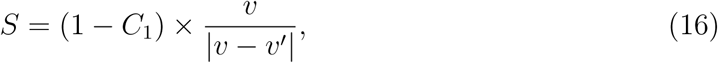

where *C*_1_ (defined in Eq. 2) ranges between 0 to 1 and measures the degree of mismatch between the calculated 3D model and the input Hi-C data, *v* is the estimated volume of the chromosome (or TAD) obtained from the prior knowledge, and *v*′ is the corresponding volume of the reconstructed model.

We use grid search to find the best pair of parameter values that yield high scores of *S* for both Chr21 and Chr22. We found that the values *λ_Ε_* = 5 × 10^12^ and *λ_F_* = 10^−8^ lead to the highest score for Chr22 and a reasonably high score for Chr21. Thus, we choose them as the default values for these two parameters when computing the TAD-level-resolution 3D models of chromosomes.

Similarly, we found that the values *λ_Ε_* = 5 × 10^11^ and *λ_R_* = 10^−7^ yield reasonably high scores of *S* for individual TADs. Thus, we choose them as the default values for these two parameters when computing the 3D conformations of individual TADs.

### 4.4 Integrating the internal conformations of TADs with the TAD-level-resolution model of the chromosome

The TAD-level-resolution 3D model of a chromosome is composed of a list of 3D points, each corresponding to a TAD. To integrate individual 3D models of TADs with this TAD-level-resolution model of the chromosome, we first translate every TAD model such that its center coincides with the corresponding point in the TAD-level-resolution model. Then, we adjust the orientation of every TAD model relative to its adjacent TADs while preserving the location of its center. This task can be achieved by rotating every TAD around its center to minimize the distance gaps between the current TAD and its adjacent ones. Reflection of a TAD model through a mirror plane passing by its center is also considered during this optimization process.

A rough estimate of the spatial distance between two adjacent TADs *i* and *i* + 1 along the genome can be derived either from the contact frequency between the last genomic locus of *TAD_i_* and the first genomic locus of *TAD_i_*_+1_, or from the relation between genomic and spatial distances in that particular chromosome if the contact frequency between these two loci is equal to zero. That is,

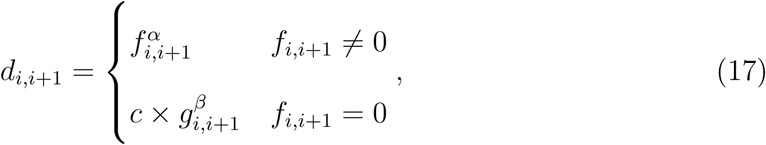

where *d_i,i_*_+1_ stands for the estimated spatial distance between the last locus of *TAD_i_* and the first locus of *TAD_i_*_+1_, *f_i_,_i_*_+1_ and *g_i,i_*_+1_ stand for the contact frequency and genomic distance between those two loci, respectively. According to the relation between spatial distance and contact frequency for a pair of TADs derived in [6], *α* is set to −0.25. The proportionality constant *c* and the scaling exponent *β* are derived from the FISH data [6] for each inspected chromosome.

The problem of placing the intra-TAD structures to the TAD-level-resolution model can be formulated as an optimization problem, with the goal to minimize the following cost function *C_integration_*,

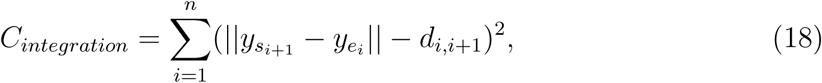

where 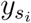 and 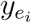 are the first and last points in the model of *TAD_i_*, respectively.

We use gradient descent to optimize *C_integration_*. In particular, the gradient of *C_integration_* with respect to 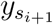 can be given by,

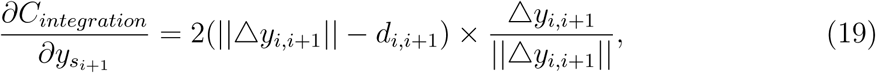

where 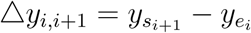.

In each iteration, until convergence of *C_integration_* is reached, 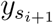 should be updated by adding a value proportional to the negative of 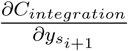. Thus, the new value of 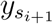, denoted by 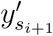, can be given by,

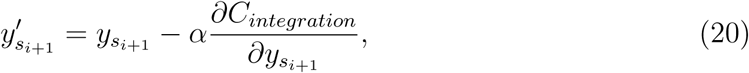

where *α* is the learning rate. However, the points associated with *TAD_i_*_+1_ are only allowed to rotate around its center. Based on the positions of 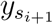, 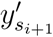, and the center of *TAD_i_*_+1_, we can force the angle and axis of rotation of the point 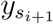 to fit into the direction towards point 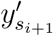. We then rotate all the points within the *TAD_i_*_+1_ model around its center using the same rotation angle.

## Data Availability

The source code of GEM-FISH can be downloaded from https://github.com/mlcb-thu/GEM-FISH. The Hi-C and FISH data of the human IMR90 cell line can be downloaded from NCBI GEO GSE63525 and https://www.sciencemag.org/content/353/6299/598/suppl/DC1, respectively.

## Acknowledgments

We thank Mohamed Ibrahim (King Abdullah University of Science and Technology) for useful discussions. Molecular graphics and analyses were performed with the UCSF Chimera package. Chimera is developed by the Resource for Biocomputing, Visualization, and Informatics at the University of California, San Francisco (supported by NIGMS P41-GM103311).

## Author Contributions

A.A., J.G., M.Z., and J.Z. conceived the research project. J.Z. supervised the research project. A.A., G.Z., and J.Z. designed the computational pipeline. A.A. implemented GEM-FISH. A.A., X.H., B.Z., Z.M., J.G., M.Z., and J.Z. discussed the modeling and validation results. A.A. and J.Z. wrote the manuscript with support from all authors.

---

**Table S1.** Consistency between the median spatial distances between the TAD pairs from FISH data [6] and their corresponding Hi-C contact frequencies [5] used in our tests, which was defined as the percentage of median spatial distances that vary concordantly with the corresponding Hi-C contact frequencies. The control maps were randomly generated with non-negative numbers, satisfying the symmetric property and having the same sizes as the Hi-C maps.

|  | Hi-C maps [5] | Control maps |
| --- | --- | --- |
| Chr20 | 79.67% | 48.72% |
| Chr21 | 83.79% | 49.13% |
| Chr22 | 82.63% | 49.78% |
| Average | 82.63% | 49.21% |

**Figure S1.**
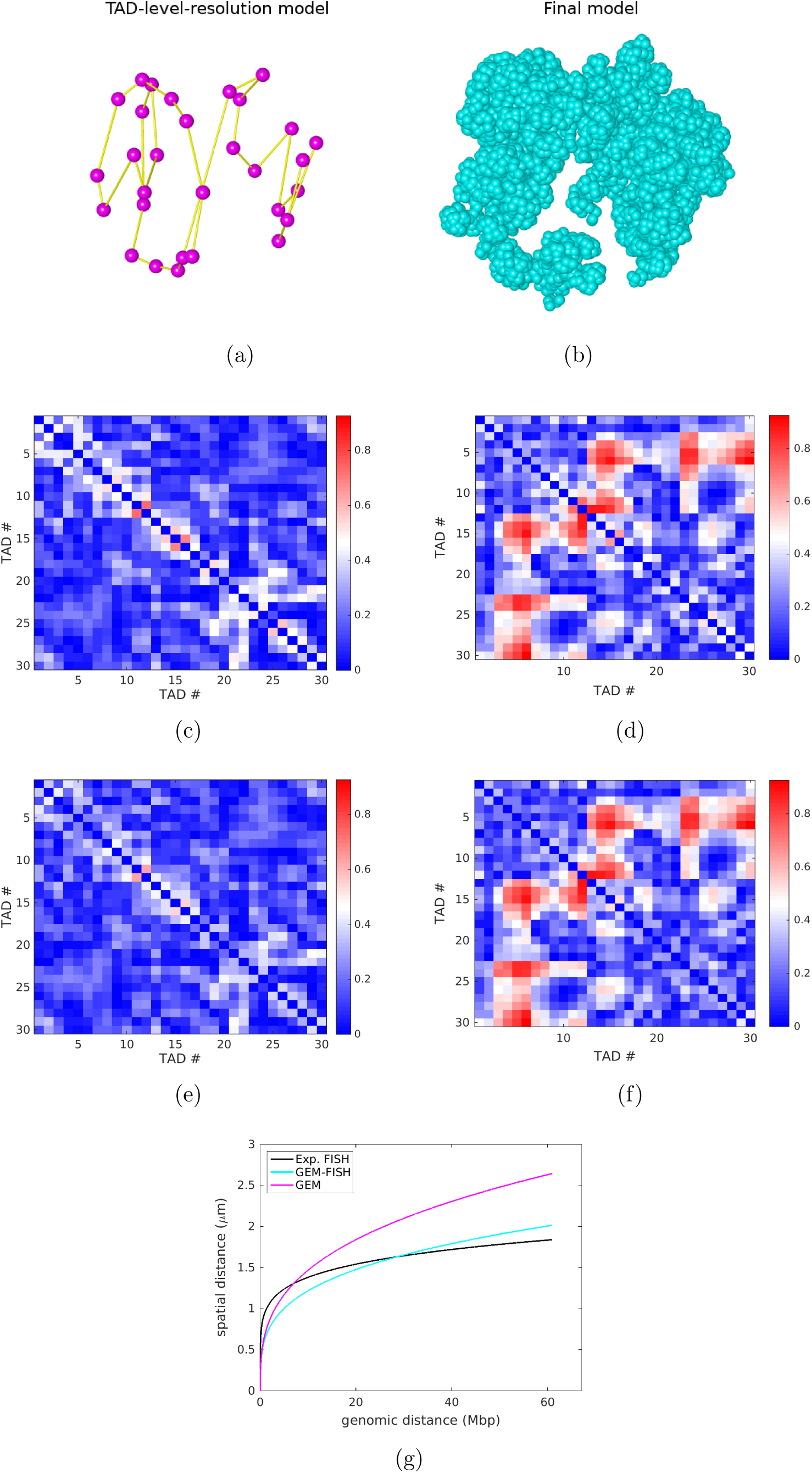
The modeling results of human Chromosome 20 (Chr20). (a) The TAD-level-resolution 3D structure of Chr20 calculated by GEM-FISH, where each dot represents the center of a TAD. (b) The final 3D structure of Chr20 reconstructed by GEM-FISH. The visualization in (a) and (b) was performed using UCSF Chimera [48]. (c and d) The relative error matrices of the TAD-level-resolution models computed by GEM-FISH using both Hi-C and FISH data, and by GEM using only Hi-C data, respectively. (e and f) The relative error matrices of the final models computed by GEM-FISH using both Hi-C and FISH data, and GEM using only Hi-C data, respectively. (g) The curves of spatial vs. genomic distances between TADs, which were derived from the experimental FISH data, and the final 3D models reconstructed by GEM-FISH (using both Hi-C and FISH data), and by GEM (using only Hi-C data), respectively.

**Figure S2.**
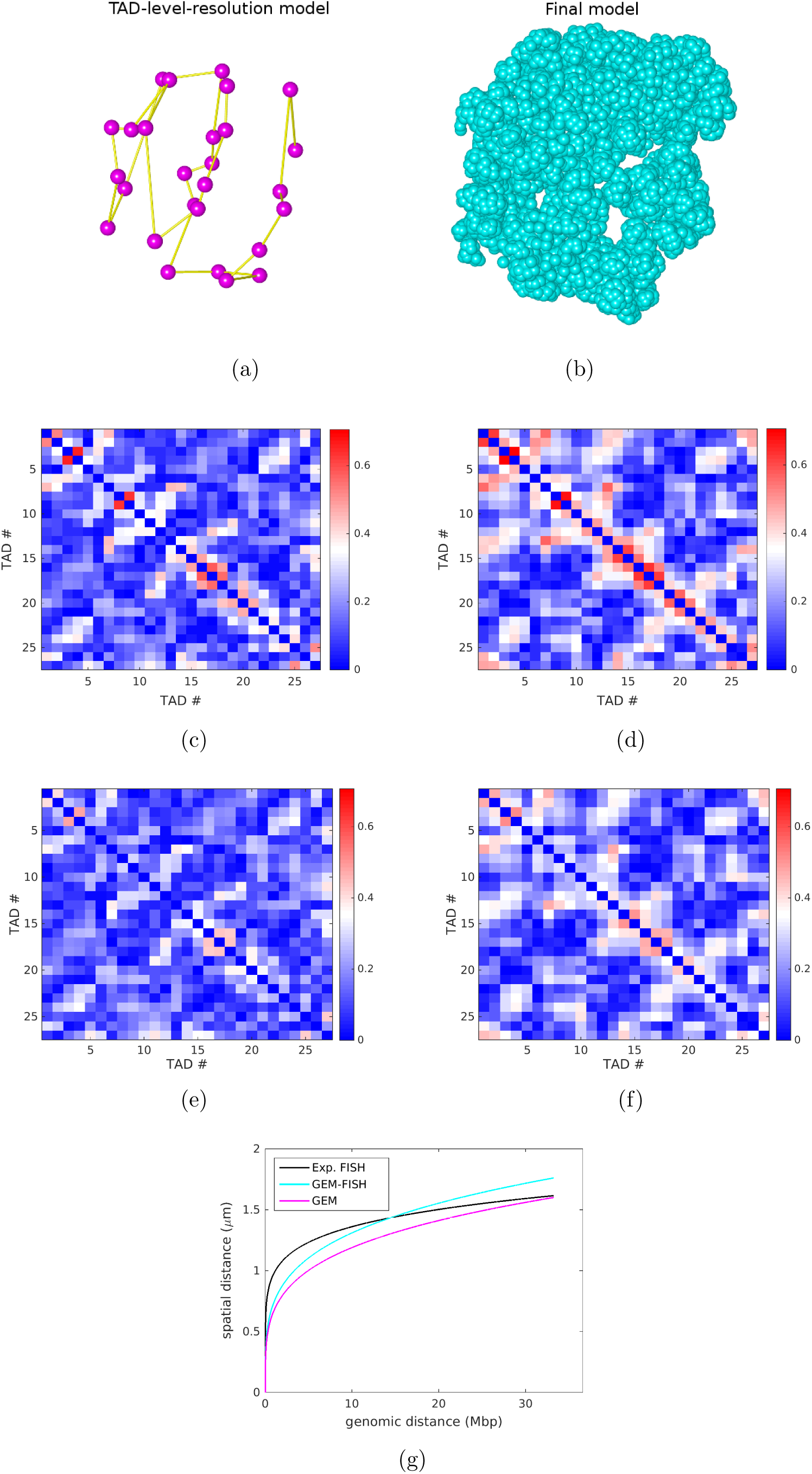
The modeling results of human Chromosome 22 (Chr22). (a) The TAD-level-resolution 3D structure of Chr22 calculated by GEM-FISH, where each dot represents the center of a TAD. (b) The final 3D structure of Chr22 reconstructed by GEM-FISH. The visualization in (a) and (b) was performed using UCSF Chimera [48]. (c and d) The relative error matrices of the TAD-level-resolution models computed by GEM-FISH using both Hi-C and FISH data, and by GEM using only Hi-C data, respectively. (e and f) The relative error matrices of the final models computed by GEM-FISH using both Hi-C and FISH data, and GEM using only Hi-C data, respectively. (g) The curves of spatial vs. genomic distances between TADs, which were derived from the experimental FISH data, and the final 3D models reconstructed by GEM-FISH (using both Hi-C and FISH data), and by GEM (using only Hi-C data), respectively.

**Figure S3.**
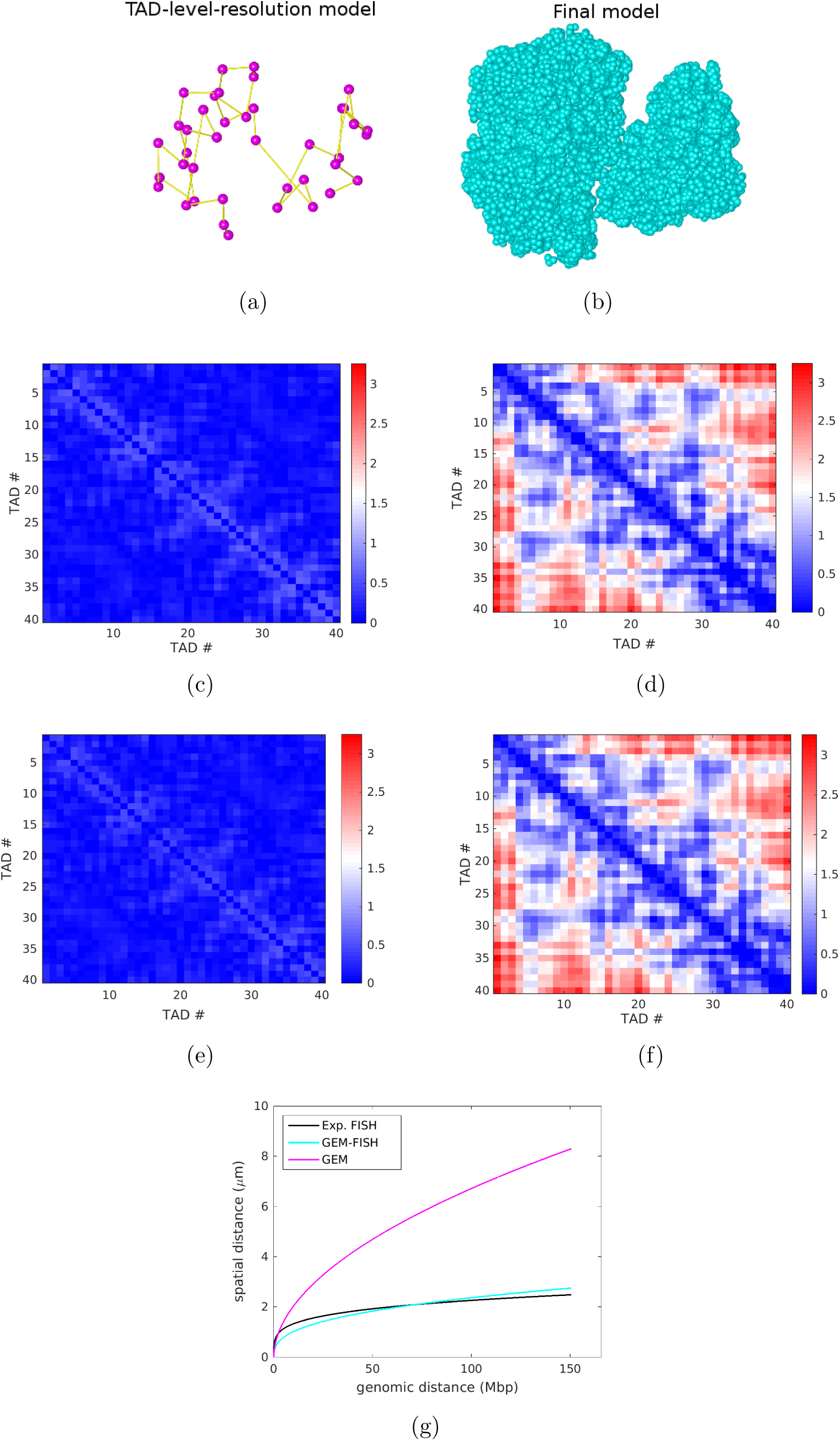
The modeling results of human Chromosome Xa (ChrXa). (a) The TAD-level-resolution 3D structure of ChrXa calculated by GEM-FISH, where each dot represents the center of a TAD. (b) The final 3D structure of ChrXa reconstructed by GEM-FISH. The visualization in (a) and (b) was performed using UCSF Chimera [48]. (c and d) The relative error matrices of the TAD-level-resolution models computed by GEM-FISH using both Hi-C and FISH data, and by GEM using only Hi-C data, respectively. (e and f) The relative error matrices of the final models computed by GEM-FISH using both Hi-C and FISH data, and GEM using only Hi-C data, respectively. (g) The curves of spatial vs. genomic distances between TADs, which were derived from the experimental FISH data, and the final 3D models reconstructed by GEM-FISH (using both Hi-C and FISH data), and by GEM (using only Hi-C data), respectively.

**Figure S4.**
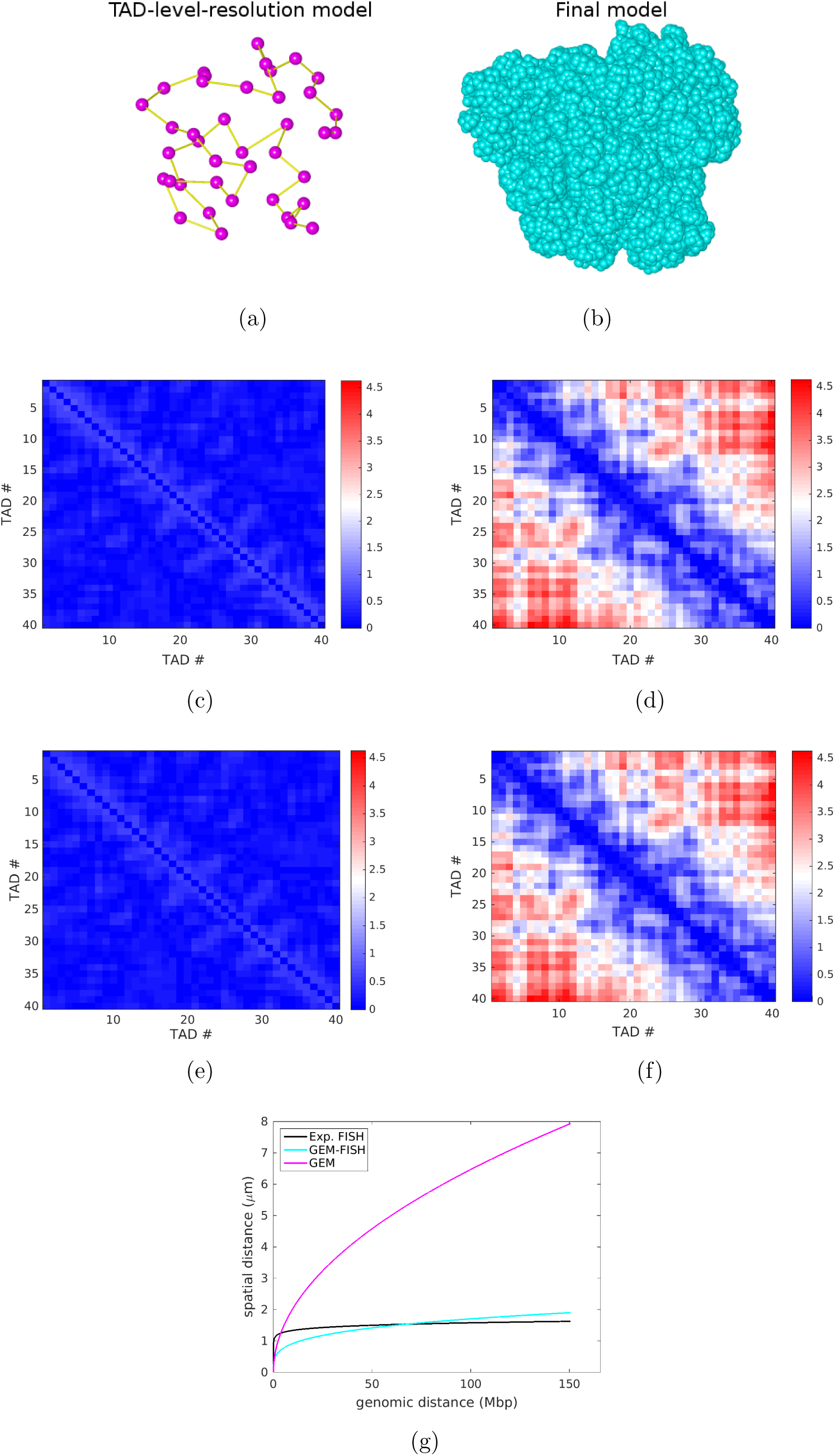
The modeling results of human Chromosome Xi (ChrXi). (a) The TAD-level-resolution 3D structure of ChrXi calculated by GEM-FISH, where each dot represents the center of a TAD. (b) The final 3D structure of ChrXi reconstructed by GEM-FISH. The visualization in (a) and (b) was performed using UCSF Chimera [48]. (c and d) The relative error matrices of the TAD-level-resolution models computed by GEM-FISH using both Hi-C and FISH data, and by GEM using only Hi-C data, respectively. (e and f) The relative error matrices of the final models computed by GEM-FISH using both Hi-C and FISH data, and GEM using only Hi-C data, respectively. (g) The curves of spatial vs. genomic distances between TADs, which were derived from the experimental FISH data, and the final 3D models reconstructed by GEM-FISH (using both Hi-C and FISH data), and by GEM (using only Hi-C data), respectively.

**Figure S5.**
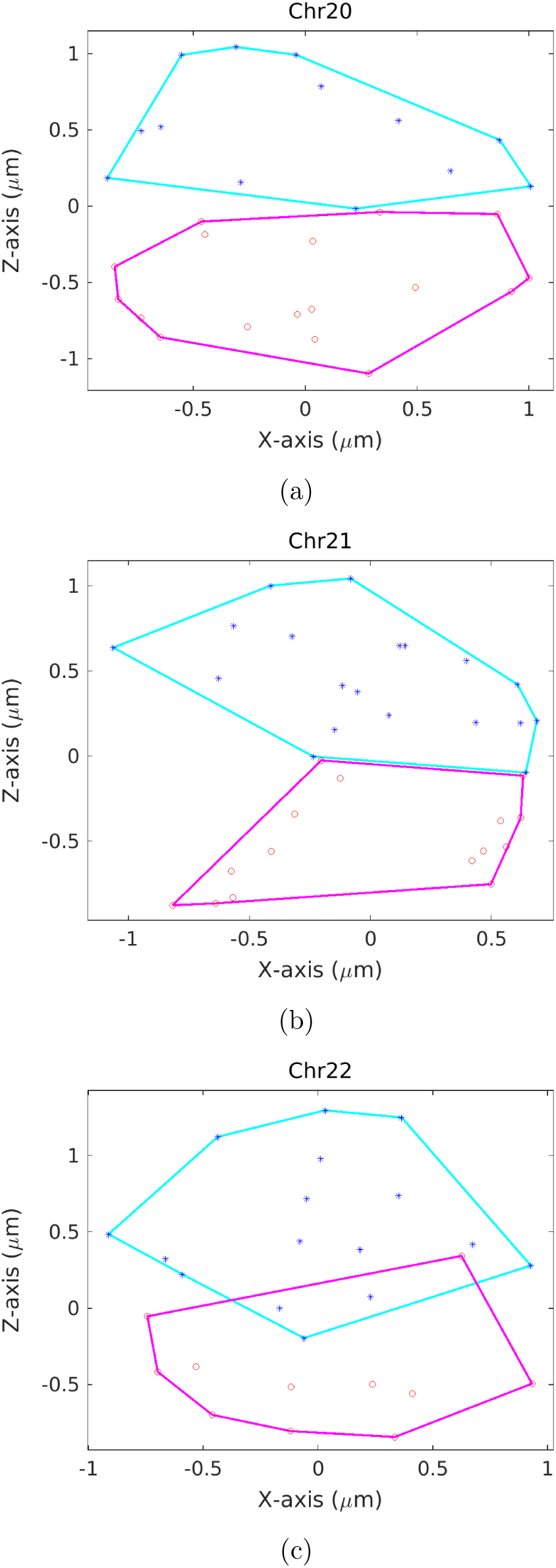
Projection of the 3D convex hull plots of the A and B compartments to the XZ plane for Chr20 (a), Chr21 (b), and Chr22 (c).

**Figure S6.**
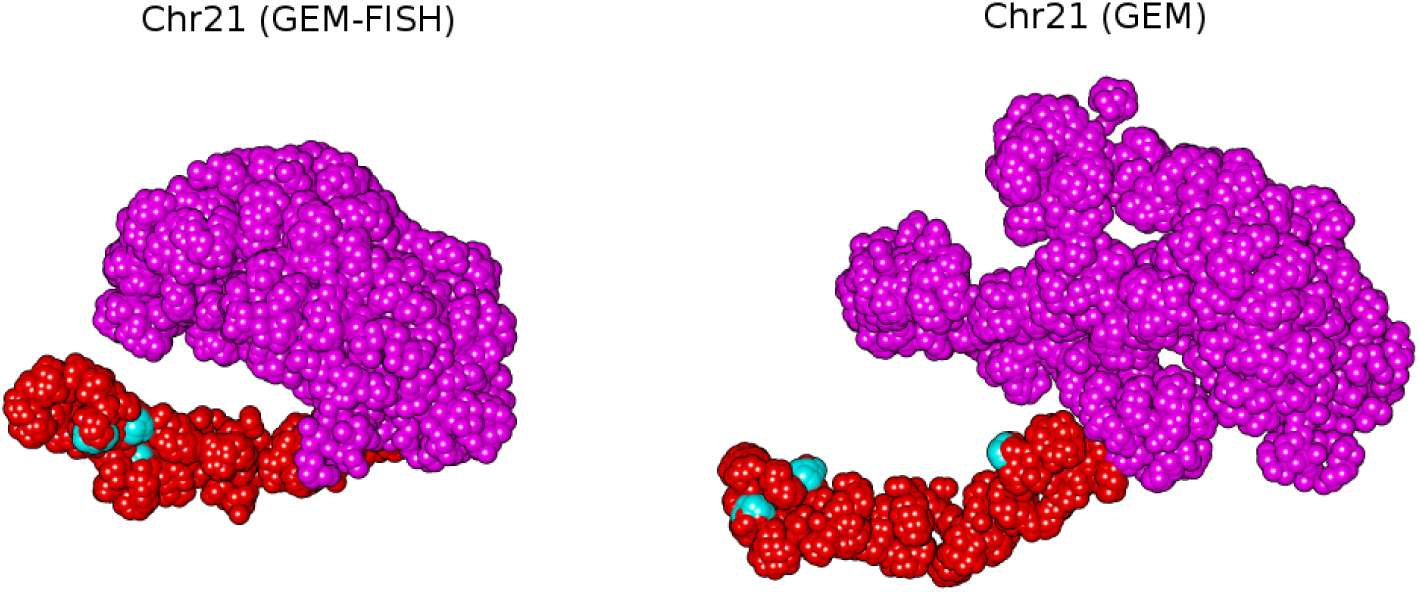
Comparison between the 3D models of Chr21 reconstructed by GEM-FISH (left) using both Hi-C and FISH data and GEM (right) using Hi-C data alone. The visualization was performed using UCSF Chimera [48]. In both 3D models, the gene-rich G-band q22.3 is shown in red, while the remaining parts are shown in magenta. The super-enhancers are shown in cyan.

**Figure S7.**
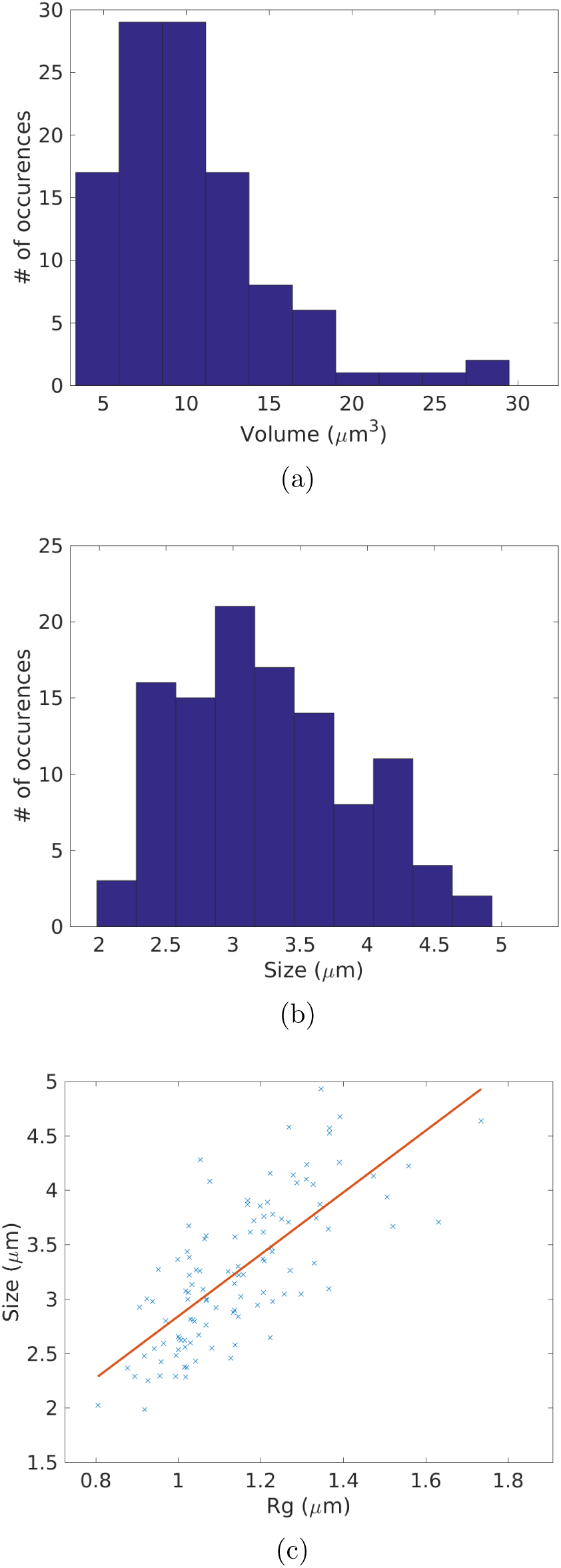
Distributions of the volumes and sizes of Chr20, and the relation between its size and its radius of gyration obtained from experimental FISH data. (a) Histogram showing the distribution of the volumes of Chr20. (b) Histogram showing the distribution of the sizes of Chr20. (c) The approximate linear relation between the radius of gyration and the size of Chr20. Data from 111 cells [6] were used to generate (a), (b), and (c).

**Figure S8.**
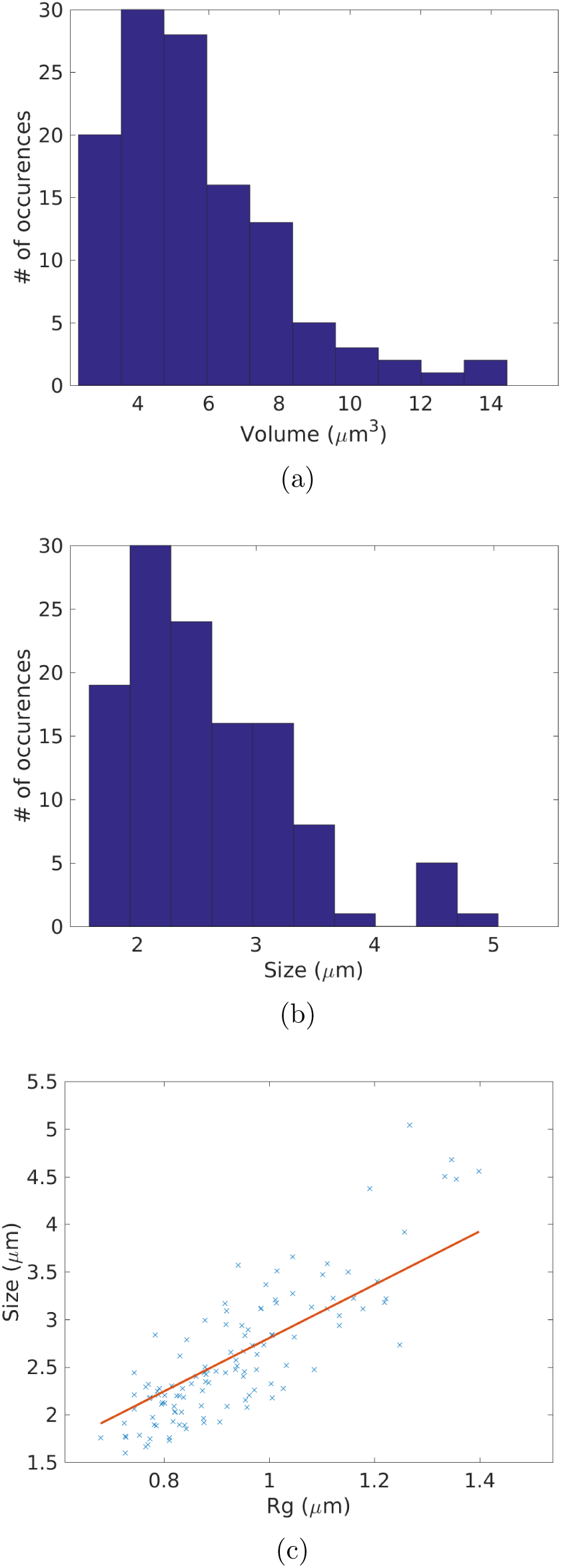
Distributions of the volumes and sizes of Chr21, and the relation between its size and its radius of gyration obtained from experimental FISH data. (a) Histogram showing the distribution of the volumes of Chr21. (b) Histogram showing the distribution of the sizes of Chr21. (c) The approximate linear relation between the radius of gyration and the size of Chr21. Data from 120 cells [6] were used to generate (a), (b), and (c).

**Figure S9.**
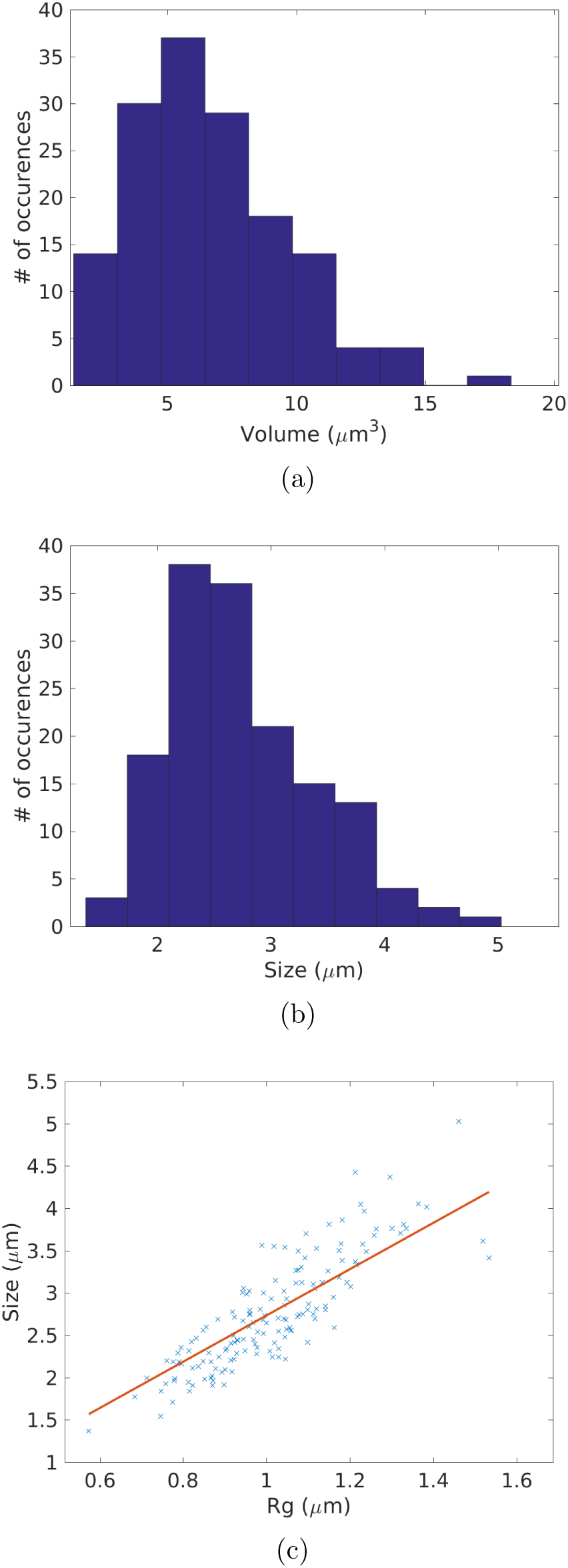
Distributions of the volumes and sizes of Chr22, and the relation between its size and its radius of gyration obtained from experimental FISH data. (a) Histogram showing the distribution of the volumes of Chr22. (b) Histogram showing the distribution of the sizes of Chr22. (c) The approximate linear relation between the radius of gyration and the size of Chr22. Data from 151 cells [6] were used to generate (a), (b), and (c).

**Figure S10.**
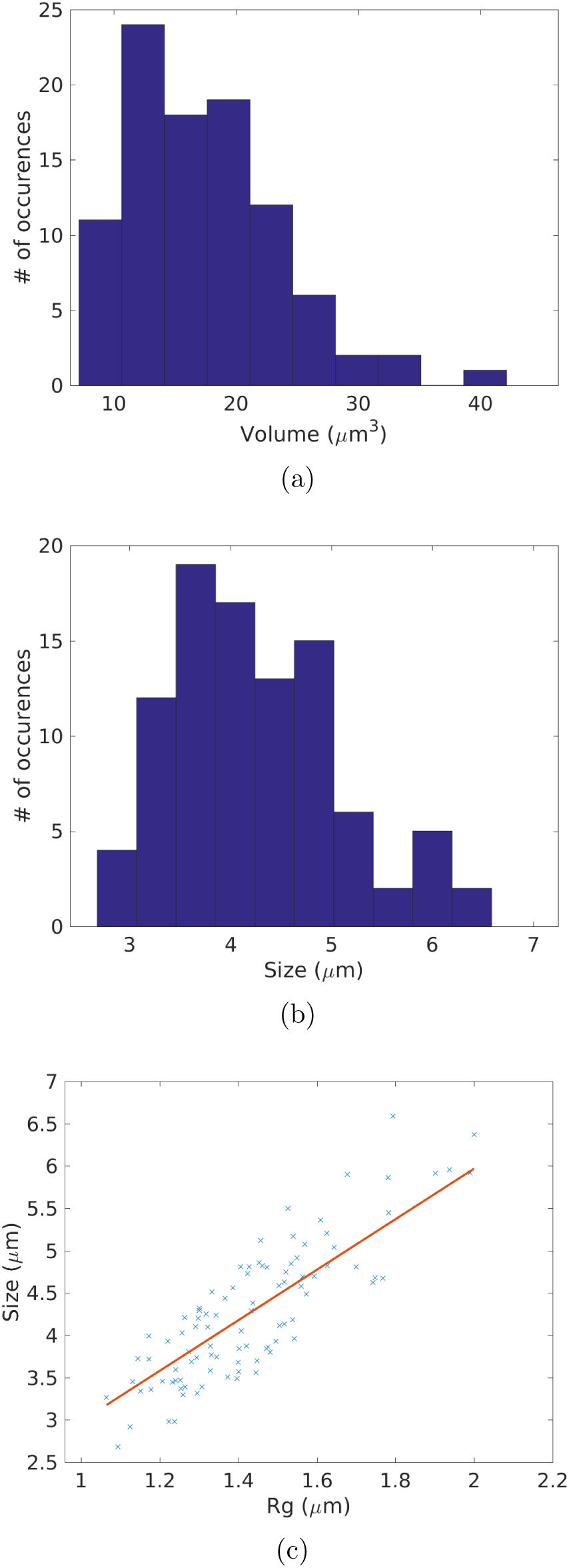
Distributions of the volumes and sizes of ChrXa, and the relation between its size and its radius of gyration obtained from experimental FISH data. (a) Histogram showing the distribution of the volumes of ChrXa. (b) Histogram showing the distribution of the sizes of ChrXa. (c) The approximate linear relation between the radius of gyration and the size of ChrXa. Data from 95 cells [6] were used to generate (a), (b), and (c).

**Figure S11.**
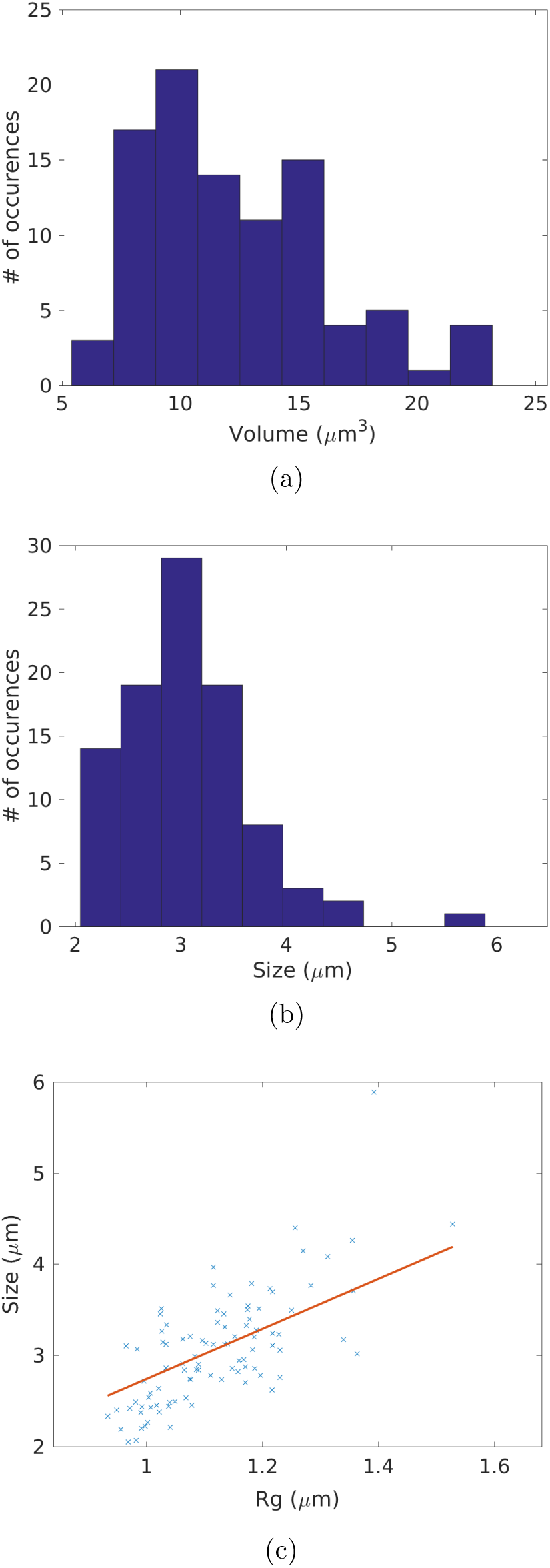
Distributions of the volumes and sizes of ChrXi, and the relation between its size and its radius of gyration obtained from experimental FISH data. (a) Histogram showing the distribution of the volumes of ChrXi. (b) Histogram showing the distribution of the sizes of ChrXi. (c) The approximate linear relation between the radius of gyration and the size of ChrXi. Data from 95 cells [6] were used to generate (a), (b), and (c).

